# Sampling from Disentangled Representations of Single-Cell Data Using Generative Adversarial Networks

**DOI:** 10.1101/2021.01.15.426872

**Authors:** Hengshi Yu, Joshua D. Welch

## Abstract

Deep generative models, including variational autoencoders (VAEs) and generative adversarial networks (GANs), have achieved remarkable successes in generating and manipulating highdimensional images. VAEs excel at learning disentangled image representations, while GANs excel at generating realistic images. Here, we systematically assess disentanglement and generation performance on single-cell gene expression data and find that these strengths and weaknesses of VAEs and GANs apply to single-cell gene expression data in a similar way. We also develop MichiGAN^1^, a novel neural network that combines the strengths of VAEs and GANs to sample from disentangled representations without sacrificing data generation quality. We learn disentangled representations of two large singlecell RNA-seq datasets [13, 68] and use MichiGAN to sample from these representations. MichiGAN allows us to manipulate semantically distinct aspects of cellular identity and predict single-cell gene expression response to drug treatment.

## 1 Introduction

Deep learning techniques have recently achieved remarkable successes, especially in vision and language applications [44,6]. In particular, state-of-the-art deep generative models can generate realistic images or sentences from low-dimensional latent variables [70]. The generated images and text data are often nearly indistinguishable from real data, and data generating performance is rapidly improving [9, 76]. The two most widely used types of deep generative models are variational autoencoders (VAEs) and generative adversarial networks (GANs). VAEs use a Bayesian approach to estimate the posterior distribution of a probabilistic encoder network, based on a combination of reconstruction error and the prior probability of the encoded distribution [41]. In contrast, the GAN framework consists of a two-player game between a generator network and a discriminator network [26]. GANs and VAEs possess complementary strengths and weaknesses: GANs generate much better samples than VAEs [25], but VAE training is much more stable and learns more useful “disentangled” latent representations [32]. Some recent GAN extensions, such as Wasserstein GAN, [2, 28, 30] significantly improve the stability of GAN training, which is particularly helpful for non-image data.

Achieving a property called “disentanglement”, in which each dimension of the latent representation controls a semantically distinct factor of variation, is a key focus of recent research on deep generative models [17,63,16,1,21,49,31]. Disentanglement is important for controlling data generation and generalizing to unseen latent variable combinations. For example, disentangled representations of image data allow prediction of intermediate images [7] and mixing images’ styles [37]. For reasons that are not fully understood, VAEs generally learn representations that are more disentangled than other approaches [33, 20, 4, 64, 23, 39]. The state-of-the-art methods for learning disentangled representations capitalize on this advantage by employing modified VAE objective functions that further improve disentanglement, including *β*-VAE, FactorVAE, and *β*-TCVAE [32,40,11,24]. In contrast, the latent space of the traditional GAN is highly entangled. Some modified GAN architectures, such as InfoGAN [12], encourage disentanglement using purely unsupervised techniques, but these approaches still do not match the disentanglement performance of VAEs [60, 36, 35, 46, 38, 66, 47, 45].

Disentanglement performance is usually quantitatively evaluated on standard image datasets with known ground truth factors of variation [52, 57, 3, 48]. In addition, disentangled representations can be qualitatively assessed by performing traversals or linear arithmetic in the latent space and visually inspecting the resulting images [10, 74, 42, 19, 65].

Recently, molecular biology has seen the rapid growth of single-cell RNA-seq technologies that can measure the expression levels of all genes across thousands to millions of cells [22]. Like image data, for which deep generative models have proven so successful, single-cell RNA-seq datasets are large and high-dimensional. Thus, it seems likely that deep learning will be helpful for single-cell data. In particular, deep generative models hold great promise for distilling semantically distinct facets of cellular identity and predicting unseen cell states.

Several papers have already applied VAEs [50, 69, 29, 73, 61, 15, 27, 72, 18, 34, 14] and GANs [51] to singlecell data. A representative VAE method is scGen, which uses the same objective function as *β*-VAE [32]. The learned latent values in scGen are utilized for out-of-sample predictions by latent space arithmetic. The cscGAN paper adapts the Wasserstein GAN approach for single-cell data and shows that it can generate realistic gene expression profiles, proposing to use it for data augmentation.

Assessing disentanglement performance of models on single-cell data is more challenging than image data, because humans cannot intuitively understand the data by looking at it as with images. Previous approaches such as scGen have implicitly used the properties of disentangled representations [50], but disentanglement performance has not been rigorously assessed on single-cell data.

In this paper, we systematically assess the disentanglement and generation performance of deep generative models on single-cell RNA-seq data. We show that the complementary strengths and weaknesses of VAEs and GANs apply to single-cell data in a similar way as image data. We also develop MichiGAN, a neural network that combines the strengths of VAEs and GANs to sample from disentangled representations without sacrificing data generation quality. We employ MichiGAN and other methods on simulated single-cell RNA-seq data [78] and provide quantitative comparisons through several disentanglement metrics [11]. We learn disentangled representations of two real single-cell RNA-seq datasets [13, 68] and show that the disentangled representations can control semantically distinct aspects of cellular identity and predict unseen combinations of cell states.

## 2 Methods

### 2.1 MichiGAN framework combines strengths of VAEs and GANs

As VAEs achieve better disentanglement performance, but GANs achieve better generation performance, we sought to develop an approach that combines the strengths of both techniques. Several previous approaches have combined variational and adversarial techniques [43, 59, 53]. However, when we tested these approaches on single-cell data, we found that attempts to jointly perform variational and adversarial training compromised both training stability and generation performance. We also investigated the InfoGAN and semi-supervised InfoGAN, but found that the disentanglement performance was still significantly worse than that of the VAE approaches.

We thus developed a different approach: we first train a VAE to learn a disentangled representation. Then, we use the VAE encoder’s latent representation *z* for each cell *x* as a given code and train a conditional GAN using the (*z, x*) pairs. After training, we can generate high-quality samples from the VAE’s disentangled representation. Importantly, the training is no less stable than training VAE and GAN separately, and the GAN generation quality is not compromised by a regularization term encouraging disentanglement. Wanting to follow the convention that the names of many generative adversarial networks end with “GAN”, but unable to devise a compelling acronym, we named our approach MichiGAN after our institution. Algorithm 1 summarizes the MichiGAN framework:

#### Algorithm 1

MichiGAN

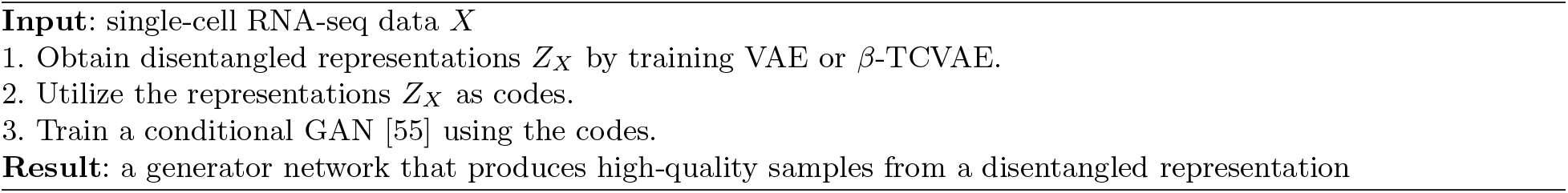

The MichiGAN framework is also shown in Fig. 1. We find that MichiGAN effectively achieves our goal of sampling from a disentangled representation without compromising generation quality (see results below). Although our approach is conceptually simple, there are several underlying reasons why it performs so well, and recognizing these led us to pursue this approach. First, training a conditional GAN maximizes mutual information between the condition variable and the generated data. This is a similar intuition as the InfoGAN, but unlike InfoGAN, MichiGAN does not need to learn its own codes, and thus the discriminator can focus exclusively on enforcing the relationship between code and data. A nearly optimal discriminator is crucial for maximizing this mutual information, but the Wasserstein loss also has this requirement, and we meet it by training the discriminator 5 times for every generator update. Second, the adversarial loss allows the GAN generator to capture complex, multi-modal distributional structure that cannot be modeled by the factorized Gaussian distribution of the VAE decoder. Thus, even though the GAN generates from the same latent representation as the VAE, the GAN can capture such “nuisance” variation through the effect of the noise variable input to the generator. Additionally, a data-dependent code (the posterior of the VAE encoder) allows the GAN to generate from a flexible latent space that reflects the data distribution, rather than an arbitrary distribution such as the commonly used standard normal. We believe this inflexibility contributes significantly to the relatively poor disentanglement performance of InfoGAN. For example, we found that InfoGAN is highly sensitive to the number and distribution chosen for the latent codes; if classes are imbalanced in the real data, it cannot learn a categorical variable that reflects the true proportions.

**Fig. 1:**
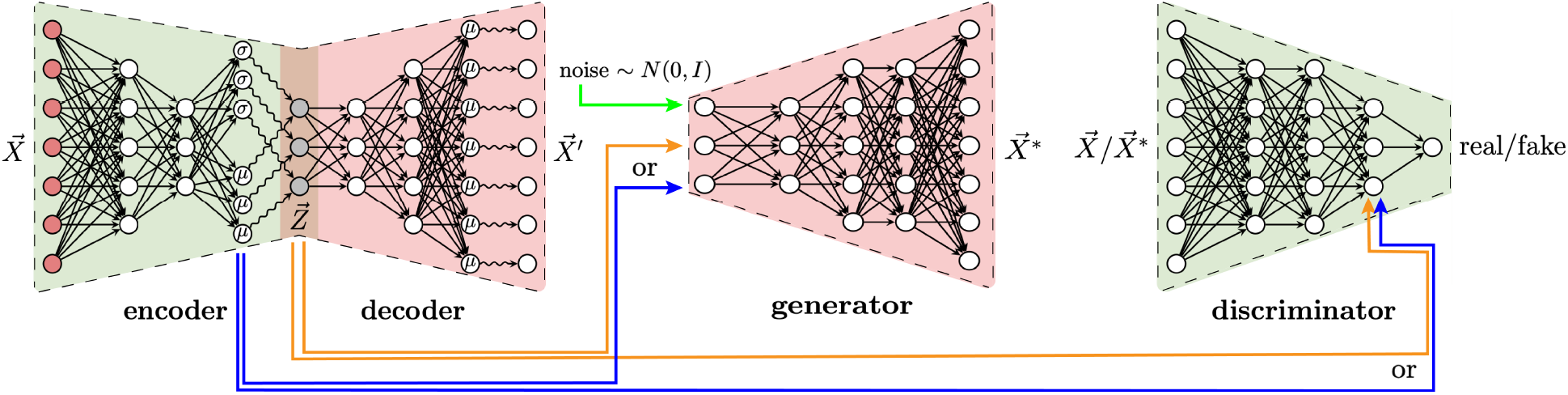
Overview of the MichiGAN Architecture.

Based on the results from our disentanglement comparison (see below), we chose to use the *β*-TCVAE to learn the latent representation for MichiGAN. We then use either the posterior means or random samples from the posterior of the VAE representation as the condition for the GAN; both choices have been utilized to evaluate disentanglement performance in previous studies [32, 40, 11].

The last step of MichiGAN involves training a conditional GAN. We found that a conditional Wasserstein GAN with projection discriminator [55] and gradient penalty [28] is most effective at enforcing the condition. We also assessed semi-supervised InfoGAN [67] and a conditional GAN based on simple concatenation, but found that these were much less effective at enforcing the relationship between code and generated data.

### 2.2 Latent space vector arithmetic

MichiGAN’s ability to sample from a disentangled representation allows predicting unseen combinations of latent variables using latent space arithmetic. We perform latent space arithmetic as in [50] to predict the single-cell gene expression of unseen cell states. Specifically, suppose we have *m* cell types *C*_1_,…, *C_m_* and *n* perturbations/treatments *D*_1_, …, *D_n_*. Denote *Z*(*C_i_*, *D_j_*) as the latent value corresponding to the expression data with combination (*C_i_, D_j_*) for 1 ≤ *i* ≤ *m* and 1 ≤ *j* ≤ *n*. If we want to predict the unobserved expression profile for the combination (*C_i′_, D_j′_*), we can calculate the average latent difference between cell type *C_i′_* and another cell type *C_k_* in the set of their joint observable treatments *Ω* that 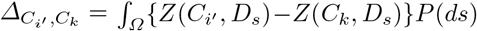 and then use the latent space *Z*((*C_k_, D_j_′*) of observed combination (*C_k_, D_j_*,) to predict

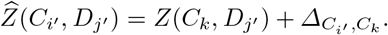

The predicted 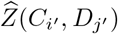 is further used to generate predicted data of the unseen combination. The predicted latent space assumes the average latent difference across jointly observed treatments is equal to the latent difference of the unobserved treatment, which may not hold if there is a strong cell type effect for the perturbation. We can test this assumption for held out data using the latent space entropy defined in the next subsection.

### 2.3 Latent space entropy

We developed a novel metric for assessing the accuracy of latent space arithmetic for a particular held-out cell type/perturbation combination. For a subset of the data *X* ~ *g*(*X*) and the latent space *Z ~ τ*(*Z*), we define the latent space entropy as:

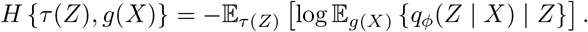

Intuitively, *H* quantifies the concentration of *Z* with respect to *X*. We can then compare the entropy of the latent embeddings for the held-out data and the latent values predicted by latent space arithmetic by calculating *ΔH* = *H*{*τ_Fake_*(*Z*), *g*(*X*)} – *H*{*τ_Real_*(*Z*), *g*(*X*)}, where *τ_Fake_* is calculated by latent space arithmetic and *τ_Real_* is calculated using the encoder. The quantity Δ*H* then gives a measure of how accurately latent space arithmetic predicts the latent values for the held-out data. If *ΔH* is positive, then the latent space prediction is less concentrated (and thus more uncertain) than the encoding of the real data.

### 2.4 Disentanglement metrics

#### Mutual information gap

Following [11], we measure the disentanglement performance of the representations using mutual information gap (MIG). The MIG metric is the difference between the largest and second largest normalized mutual information values for each ground-truth variable, averaged across all ground-truth variables. The MIG value thus indicates how much each ground truth variable is represented by exactly one dimension of the latent space. If the averaged gap is high, more mutual information is contained in one specific representation for each ground-truth variable. As described in [11], the MIG metric has the axis-alignment property, unlike the FactorVAE metric, and is unbiased for all hyperparameter settings.

#### FactorVAE metric

For completeness, we also calculated the disentanglement metric introduced in the FactorVAE paper [40]. In each of multiple repetitions, we first randomly choose a ground-truth variable and then generate data, keeping this variable fixed and other variables at random. We normalize each dimension by the empirical standard deviation over the whole data and choose the dimension with the lowest empirical variance. The pair of the dimension with the lowest empirical variance and the fixed ground-truth variable are then used to train a majority vote classifier. The disentanglement metric is then defined as the accuracy of the resulting classifier.

#### Spearman correlation gap

Inspired by the MIG metric, we also utilized Spearman correlation to quantify disentanglement performance. Although Spearman correlation is a more restricted metric of statistical dependence than mutual information, it has the advantage that it can be computed without a distributional estimate of a latent representation, which is not available for GAN models. Given the Spearman correlation *S* = cor(*Z_j_*, *V_k_*) between inferred representation *Z_j_* and ground truth variable *V_k_*, we define the corresponding Spearman correlation gap as |cor(*Z*_*j*^(*k*)^_, *V_k_*)| – max_*j*≠*j*^(*k*)^_ |cor(*Z_j_*, *V_k_*)|, where *j*^(*k*)^ = arg max_*j*_ |cor(*Z_j_*, *V_k_*)|.

### 2.5 Generation metrics

#### Random forest error

We follow the random forest error metric introduced in the cscGAN paper [51] to quantify how difficult it is for a random forest classifier to distinguish generated cells from real cells. A higher random forest error indicates that the generated samples are more realistic. We randomly sample 3,000 cells, generate 3,000 fake cells and train a random forest classifier on the 50 principal components of the 6,000 cells to predict each cell to be a real or fake cell. The random forest error is then defined as the average prediction error from 5-fold cross validation on the test data.

#### Inception score

We also define an inception score metric similar to the one widely-used in image data [5]. Intuitively, to achieve a high inception score, a generative model must generate every class in the training dataset and every generated example must be recognizable as belonging to a particular class. We train a random forest classifier on 3,000 randomly sampled real cells to predict their cell types. Based on the trained cell-type classifier, we are able to predict the probabilities of being different cell types for each generated cell. We then input the predicted probabilities to the calculation of the inception score exactly as with image data.

### 2.6 Simulating single-cell data with ground truth variables to measure disentanglement

Real single-cell datasets usually have unknown, unbalanced, and complex ground-truth variables, and humans cannot readily distinguish single-cell expression profiles by eye, making it difficult to assess disentanglement performance by either qualitative or quantitative evaluations. We thus first performed simulation experiments to generate balanced single-cell data with several data generating variables using the Splatter R package [78]. All the datasets were processed using the SCANPY software [75]. We measured the disentanglement performances of different methods on the simulated single-cell data using several disentanglement metrics and also provided qualitative evaluations on the learned representations using the real datasets.

### 2.7 Related work

To our knowledge, no approach like MichiGAN has been published. Several previous approaches have combined variational and adversarial techniques, including the VAEGAN [43], adversarial symmetric variational autoencoder [59], and adversarial variational Bayes [53]. InfoGAN and semi-supervised InfoGAN are also conceptually related to MichiGAN, but we found that none of these previously approaches produced good results on single-cell data. While we were writing this paper, another group released a preprint with an approach called ID-GAN, which also uses a pre-trained VAE to learn a disentangled representation [45]. However, they use the reverse KL divergence framework to enforce mutual information between the VAE representation and the generated data, which we previously tested and found does work as well as a conditional GAN with projection discriminator [55]. Furthermore, ID-GAN uses a convolutional architecture and classic GAN loss for image data, whereas we use a multilayer perceptron architecture and Wasserstein loss for single-cell expression data.

## 3 Results

### 3.1 Variational autoencoders learn disentangled representations of single-cell data

We first estimated simulation parameters to match the Tabula Muris dataset [13]. Then, we set the differential expression probability, factor location, factor scale, and common biological coefficient of variation to be (0.5, 0.01, 0.5, 0.1). We then used Splatter [78] to simulate gene expression data of 10, 000 cells with four underlying ground-truth variables: batch, path, step, and library size. Batch is a categorical variable that simulates differences among biological or technical replicates. Step represents the degree of progression through a simulated differentiation process, and path represents different branches of the differentiation process. We simulated two batches, two paths, and 20 steps. The batch and path variables have linear effects on the simulated expression data, while the step variable can be related either linearly or non-linearly to the simulated gene expression values. We tested the effects of this variable by separately simulating a purely linear and a non-linear differentiation process. We also included library size, the total number of expressed mRNAs per cell, as a ground truth variable. A UMAP plot of the simulated data shows that the four ground truth variables each have complementary and distinct effects on the resulting simulated gene expression state (Fig. 2a and Fig. S2a).

**Fig. 2:**
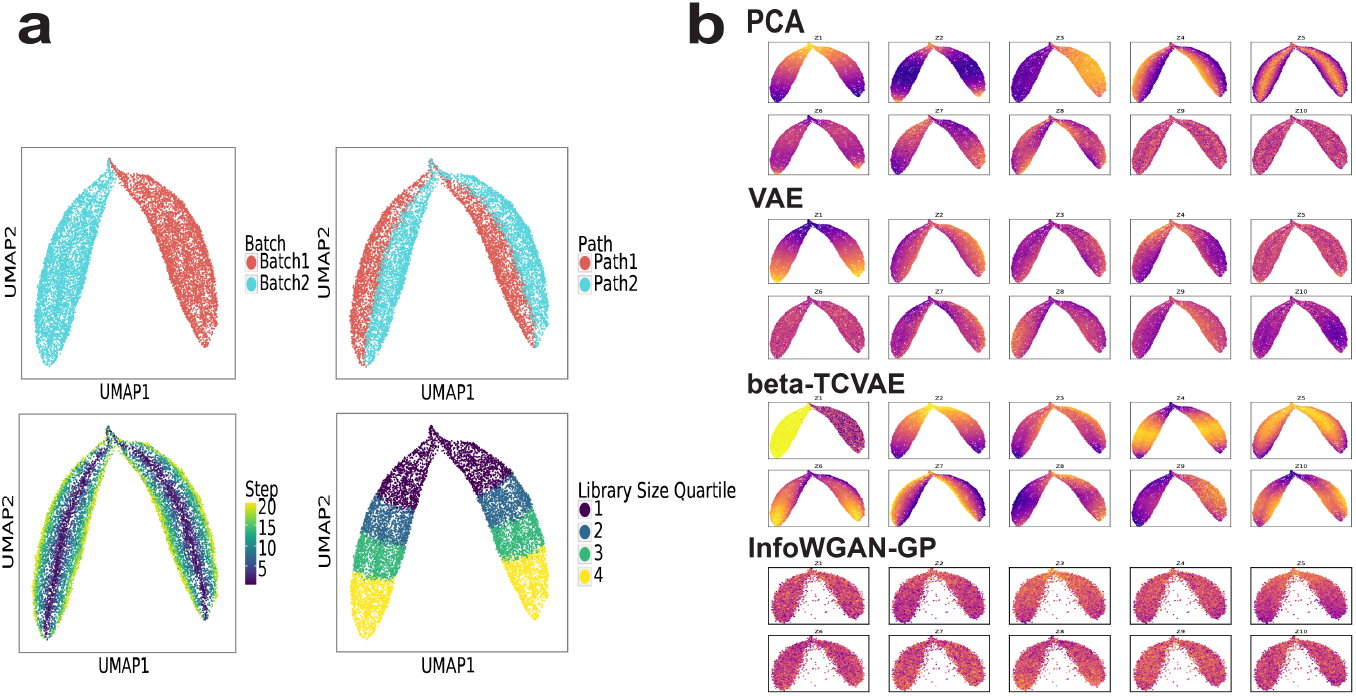
Learned representations of the simulated single-cell data with non-linear step. **a** UMAP plots of data by batch, path, step and library size quartile. **b** UMAP plots of data colored by the ten representations learned by PCA, VAE, *β*-TCVAE and InfoWGAN-GP.

We compared the disentanglement performance of three learning methods: probabilistic principal component analysis (PCA) [71], VAE, *β*-TCVAE and InfoWGAN-GP. The probabilistic PCA method assumes a linear relationship between data and representations, while VAE, *β*-TCVAE and InfoWGAN-GP can learn nonlinear representations. Note that we use probabilistic PCA to allow calculation of mutual information (see below). The *β*-TCVAE approach penalizes the total correlation of the latent representation, directly minimizing the mutual information between latent dimensions, which has been shown to significantly improve disentanglement performance on image data. The InfoWGAN-GP attempts to learn a disentangled representation by maximizing the mutual information between the latent code and generated data.

We used the four methods to learn a 10-dimensional latent representation of the simulated data (Fig. 2b, Fig. S2b and Fig. S3a). Some latent variables learned by PCA, VAE and *β*-TCVAE showed clear relationships with the ground-truth variables. For example, the first latent variable Z1 from PCA seems related to library size, and Z3, Z4, and Z5 are related to batch, path and step, respectively. The VAE representations similarly showed some relationships with the ground-truth variables. Based on the UMAP plots, the latent variables from *β*-TCVAE appear to show the strongest and most clear relationships with the ground-truth variables. In contrast, the representations of InfoWGAN-GP are entangled, and show no clear relationship to the ground-truth variables.

To quantify the disentanglement performances of the three methods of PCA, VAE and *β*-TCVAE, we calculated Spearman correlation, normalized mutual information between each representation and a groundtruth variable (Fig. S1a-b). Spearman correlation measures the strength of monotonic relatedness between two random variables. The normalized mutual information, on the other hand, is a more general and robust metric of statistical dependence. A disentangled representation should have a bar plot with only four distinct bars in this case, indicating that each ground-truth variable was captured by exactly one latent variable. PCA showed the best performance as measured by Spearman correlation (Fig. S1a), likely because the metric does not fully characterize the statistical dependency between true and inferred latent variables for the VAE methods, which learn more complex nonlinear relationships. Based on the normalized mutual information metric, both the PCA and VAE representations achieved some degree of disentanglement, but neither approach fully disentangled all ground-truth variables. Multiple PCA representations had measurable mutual information with step and library size quartile, while multiple VAE representations identified batch and path and none of the VAE representations identified step. In contrast, exactly one *β*-TCVAE representation had significant mutual information for each ground-truth variable. Also, *β*-TCVAE was the only method with a unique representation for the non-linear step variable.

We also computed the Spearman correlation and normalized mutual information for the simulated data with linear step in (Fig. S2c and Fig. S2d). The results for the simulated data with linear step were similar and *β*-TCVAE did the best at identifying only one representation for each ground-truth variable. In Fig. S3c-d, we also present Spearman correlation results for InfoWGAN-GP representations and the representations are entangled with overall small correlations with the ground-truth variables. Note that we do not report FactorVAE or MIG metrics for the InfoWGAN-GP because there is no way to calculate mutual information for GAN models, and thus we cannot calculate these metrics.

We further calculated the FactorVAE metric [40] and mutual information gap (MIG) metric used in [11] to measure disentanglement. We also calculated a similar Spearman correlation gap. Table 1 summarizes the FactorVAE metric, correlation gap and MIG of the four models for the two simulated datasets. As expected from the bar charts, the PCA representations have the largest Spearman correlation gap and *β*-TCVAE has the largest MIG, showing the best disentanglement performance for both simulated datasets. In addition, the FactorVAE metric shows that *β*-TCVAE has the best disentanglement performance for both datasets.

**Table 1:** Disentanglement metrics for two simulated single-cell RNA-seq datasets based on four ground truth variables (step, batch, path and library size)

| | | FactorVAE metric ↑ | Spearman correlation gap ↑ | Mutual information gap (MIG) ↑ |
| --- | --- | --- | --- | --- |
| Linear step | PCA | 0.36 | **0.68** | 0.53 |
|  | VAE | 0.39 | 0.43 | 0.53 |
| | β-TCVAE | **0.44** | 0.25 | **0.72** |
|  | InfoWGAN-GP | — | 0.08 | — |
| Non-linear step | PCA | 0.35 | **0.72** | 0.55 |
|  | VAE | 0.43 | 0.24 | 0.37 |
| | β-TCVAE | **0.46** | 0.17 | **0.67** |
|  | InfoWGAN-GP | — | 0.07 | — |

In summary, our assessment indicates that *β*-TCVAE most accurately disentangles the latent variables underlying single-cell data, which is consistent with its previously reported superior disentanglement performance on image data [11].

### 3.2 GANs generate more realistic single-cell expression profiles than VAEs

We next evaluated the data generating performance of several deep generative models including VAE, *β*-TCVAE and Wasserstein GAN with gradient penalty (WGAN-GP) on the Tabula Muris dataset[13]. This dataset represents a comprehensive collection of single-cell gene expression data from nearly all mouse tissues, and thus represents an appropriate dataset for evaluating data generation, analogous to the ImageNet dataset in computer vision. We also measured data generation performance on a subset of the Tabula Muris containing only cells from the mouse heart. We used two metrics to assess data generation performance: random forest error and inception score.

We show the random forest errors of VAE, *β*-TCVAE and WGAN-GP during training for the Tabula Muris heart subset and the whole Tabula Muris in Fig. S4a and Fig. S4b. WGAN-GP achieves the best generation performance, as measured by both metrics, on both the subset and full dataset. VAE achieves second-best generating performance and, as expected with an endeavour to pursue more disentangled representation, the quality of *β*-TCVAE generation is the worst of the three approaches. Fig. S4c and Fig. S4d show the inception scores for the two datasets and have consistent patterns to the random forest errors, indicating that WGAN-GP has the best generation performance and *β*-TCVAE generates the least realistic data. The random forest errors and inception scores of the three methods after training are also presented in Table 2. These results accord well with previous results from the image literature, indicating that GANs generate better samples than VAEs, and VAE modifications to encourage disentanglement come at the cost of sample quality. Interestingly, the random forest error gap between GANs and VAEs is larger on the Tabula Muris heart subset. This suggests that the adversarial training process may be able to better utilize smaller datasets.

**Table 2:** Generation metrics for the Tabula Muris heart data and the whole Tabula Muris data

| | | Random forest error (%) ↑ | Inception score ↑ |
| --- | --- | --- | --- |
| Tabula Muris heart data | VAE | 17.2 | 1.58 |
| | β-TCVAE | 10.5 | 1.49 |
|  | WGAN-GP | **24.2** | **2.01** |
| The whole Tabula Muris data | VAE | 18.1 | 3.03 |
| | β-TCVAE | 14.8 | 2.76 |
|  | WGAN-GP | **23.3** | **4.39** |
|  | MichiGAN | 21.6 | 2.69 |

### 3.3 MichiGAN samples from disentangled representations without sacrificing generation performance

We evaluated the MichiGAN algorithm on the simulated single-cell data with the trained *β*-TCVAE models. Fig. S5a shows the UMAP plots of generated data colored by the latent representations using WGAN-GP, *β*-TCVAE and MichiGAN on the simulated data with non-linear step. The WGAN-GP representations are very entangled and none of the representations shows an identifiable coloring pattern from its value. In contrast, the UMAP plots have consistent coloring patterns between the *β*-TCVAE and MichiGAN representations. Thus, the generator of MichiGAN preserves the relationship between latent code and generated data, effectively sampling from the disentangled representation learned by the *β*-TCVAE. As there is no inference network for the generated data of either WGAN-GP or MichiGAN, we are unable to measure the mutual information for the generators. Therefore, we used Spearman correlation as an indicator of whether the generated data by WGAN-GP or MichiGAN retains the disentangled representations for the variables. Fig. S5b also shows the bar plots of Spearman correlations between representations and variables for the three methods. We used the correlations between each representation and ground truth variables for *β*-TCVAE. For WGAN-GP and MichiGAN, we trained a k-nearest neighbors regressor (*k* = 3) for each variable based on the real data and predicted the variables for the generated data. As can be seen, the WGAN-GP representations do not show large correlation to any inferred ground-truth variable. In contrast, the representations for *β*-TCVAE and MichiGAN show nearly identical correlations to the true variables in the real data and predicted variables in the generated data, respectively.

We present the UMAP plots colored by the representations as well as bar plots of correlations for the simulated data with linear step in Fig. S6a and Fig. S6b. The results for the simulated data with linear step also indicate that MichiGAN restores the disentanglement performance of *β*-TCVAE, while the WGAN-GP representations are entangled. We further summarize the correlation gaps for the three methods on two simulated data in Table 3. For each simulated dataset, the MichiGAN and *β*-TCVAE have very similar correlation gaps and WGAN-GP has small correlation gap as expected.

**Table 3:** Spearman correlation gap for WGAN-GP, InfoWGAN-GP, *β*-TCVAE and MichiGAN on the two simulated single-cell RNA-seq datasets

| | WGAN-GP | InfoWGAN-GP | β-TCVAE | MichiGAN |
| --- | --- | --- | --- | --- |
| Linear step | 0.13 | 0.08 | 0.25 | 0.22 |
| Non-linear step | 0.07 | 0.07 | 0.17 | 0.15 |

We evaluated MichiGAN on the whole Tabula Muris data in Fig. S5c. Table 2 also shows the random forest errors and inception scores for the the previous three models VAE, *β*-TCVAE, WGAN-GP as well as MichiGAN after training. MichiGAN greatly improves the data generation performance based on the disentangled representations of *β*-TCVAE. Although its inception score is close to that of *β*-TCVAE, MichiGAN’s random forest error is larger than VAE and nearly as good as the WGAN-GP, while still generating samples from a disentangled latent space.

### 3.4 MichiGAN enables semantically meaningful latent traversals

Disentangled representations of images are often evaluated qualitatively by performing latent traversals, in which a single latent variable is changed holding the others fixed. Looking at the resulting changes in the generated images to see whether only a single semantic attribute changes provides a way of visually judging the quality of disentanglement. We wanted to perform a similar assessment of MichiGAN, but single-cell gene expression values are not individually and visually interpretable in the same way that images are. We thus devised a way of using UMAP plots to visualize latent traversals on single-cell data.

We perform latent traversals using both the Tabula Muris dataset and data from the recently published sci-Plex protocol [68]. After training on the Tabula Muris dataset (Fig. S7a), we chose a starting cell type, cardiac fibroblasts (Fig. S7b). We then varied the value of each latent variable from low to high, keeping the values of the other variables fixed to the latent embedding of a particular cell. For the sci-Plex dataset, which contains single-cell RNA-seq data from cells of three types (A549, K562, MCF7; Fig. S8a) and treated with one of 188 drugs, we subsampled the data to include one drug treatment from each of 18 pathways by selecting the drug with the largest number of cells (Fig. S8b). This gives us one treatment for each pathway; the numbers of cells for each combination are shown in Table S1. We then performed latent traversals on cells with cell type MCF7 and treatment S7259 (Fig. S8c).

To visualize the traversals, we plotted each of the generated cells on a UMAP plot containing all of the real cells and colored each generated cell by the value of the latent variable used to generate it. Fig. 3a-b show how traversing the latent variables concentrates the generated values on each part of the UMAP plots for Tabula Muris data using the first 10 dimensions of 128-dimensional WGAN-GP and MichiGAN, respectively. Figure 3c-d are the latent-traversal plots for the sci-Plex data using WGAN-GP and MichiGAN. As shown in Fig. 3b, all but three of the latent variables learned by the *β*-TCVAE behave like noise when we traverse them starting from the fibroblast cells, a property previously noted in assessments of disentangled latent variables learned by VAEs [40]. The remaining dimensions, Z3, Z6, and Z10, show semantically meaningful latent traversals. Latent variable Z3 shows high values for mesenchymal stem cells and fibroblasts, with a gradual transition to differentiated epithelial cell types from bladder, intestine, and pancreas at lower values of Z3. This is intriguing, because the mesenchymal-epithelial transition is a key biological process in normal development, wound healing, and cell reprogramming [58]. Latent variable Z6 generates mesenchymal and endothelial cells at low values, and mammary epithelial and cardiac muscle cells at high values. Latent variable Z10 is clearly related to immune function, generating immune cells at low and medium values and traversing from hematopoietic stem and progenitor cells to monocytes, T cells, and B cells. In contrast, latent traversals in the latent space of 128-dimensional WGAN-GP (Fig. 3a) do not show semantically meaningful changes along each dimension.

**Fig. 3:**
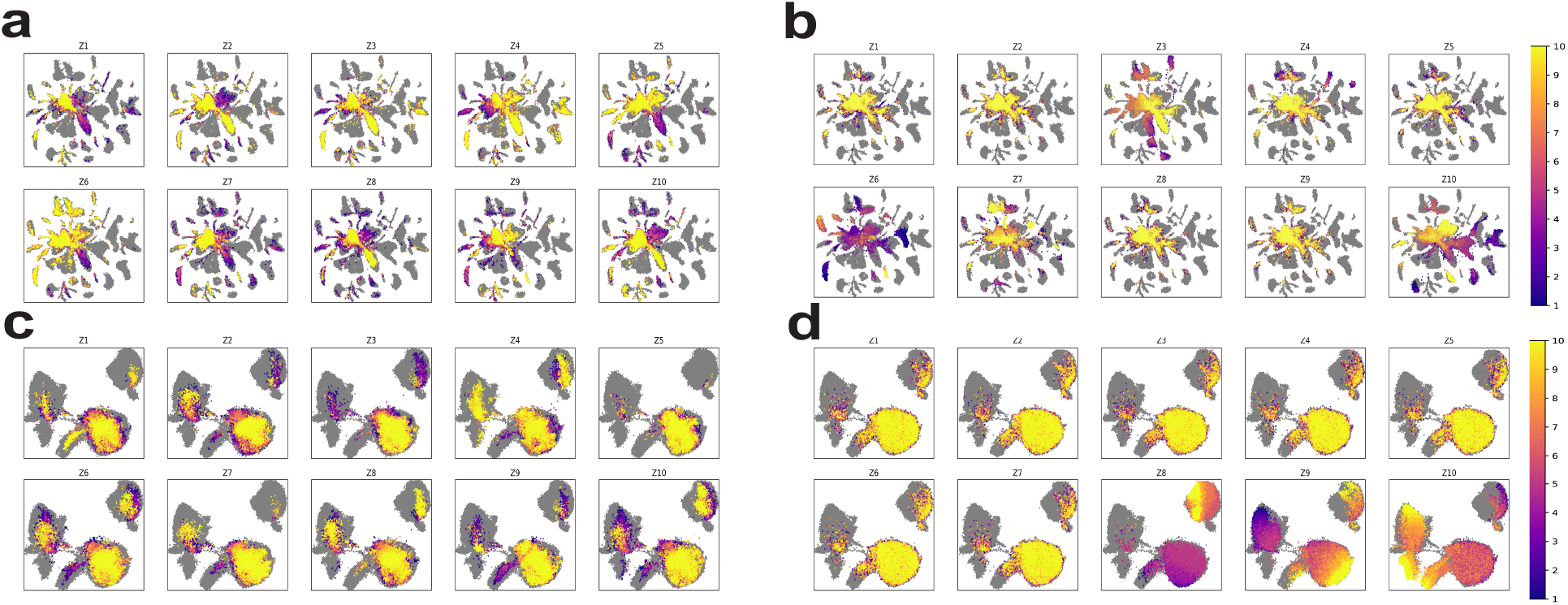
Latent traversals of WGAN-GP and MichiGAN on Tabula Muris and sci-Plex datasets. **a** UMAP plot of latent traversals of the 10 representations of latent values that generate data closest to fibroblast cells in heart within the Tabula Muris data using WGAN-GP with 128 dimensions. **b** UMAP plot of latent traversals of the 10 representations of latent values of fibroblast cells in heart within the Tabula Muris data using MichiGAN. **c** UMAP plot of latent traversals of the 10 representations of latent values that generate data closest to MCF7-S7259 cells within the sci-Plex data using WGAN-GP with 128 dimensions. **d** UMAP plot of latent traversals of the 10 representations of latent values of MCF7-S7259 cells within the sci-Plex data using MichiGAN.

Fig. 3d also shows that MichiGAN’s latent traversals gives meaningful changes on the sci-Plex data. Latent variable Z8 has lower values on MCF7 cells and gradually transitions to higher values on K562 cells. In addition, latent variable Z9 also shows an A549-MCF7 transition with lower values on the A549 cells. The latent traversals of the 128-dimensional WGAN-GP, however, do not provide interpretable changes across the UMAP plot along each dimension. We also provide the latent traversals using 10-dimensional WGAN-GP for the two datasets in Fig. S9a-b and find that the latent traversals are still not semantically meaningful.

### 3.5 MichiGAN predicts single-cell gene expression changes under unseen drug treatments

One of the most exciting applications of disentangled representations is predicting high-dimensional data from unseen combinations of latent variables. We next investigated whether MichiGAN can predict singlecell gene expression response to drug treatment for unseen combinations of cell type and drug. We note that this is a *data generation* task that cannot be solved by straightforward application of supervised learning techniques such as logistic regression, random forests, or feedforward neural networks. Such approaches have been applied to predict the mean expression of *bulk samples* under different drug treatments, but cannot be used to generate samples from the distribution of possible single-cell gene expression states.

We trained MichiGAN on the subsampled sci-Plex data [68] with three cell types (A549, K562, MCF7) and 18 drug treatments. We also held out three drug/cell type combinations (A549-S1628, K562-S1096 and MCF7-S7259) to test MichiGAN’s out-of-sample prediction ability. We predict single-cell gene expression for each drug/cell type combination in a two-step process. First, we estimate the mean latent difference between the target cell type and another control cell type for other treatments using either posterior means or posterior samples from the *β*-TCVAE encoder. We then add the average latent difference to the latent values with the same treatment and the control cell type. This latent space vector arithmetic assumes the mean cell type latent differences are homogeneous across different treatments. Note that this assumption may not hold if there is a strong interaction effect between cell type and drug treatment.

Because there are a total of three cell types, we have a total of six predictions for the three held-out drug/cell type combinations. Fig. S10a shows UMAP plots for these six predictions. For all six predictions, the predicted values are closer to the true drug treated cells on the UMAP plot than the control cells used to calculate the latent vector. However, the predicted cells do not overlap with the treated cells for the combinations A549-S1628 and K562-S1096, while the two predictions for MCF7-S7259 appear to be more accurate. For both *β*-TCVAE and MichiGAN, we measure their random forest errors between the real and predicted cells for each combination. The random forest error scatter plots for sampled representations are shown in Fig. 4a. The MichiGAN framework with sampled representations has significantly better random forest errors than *β*-TCVAE (*p* < 10^−4^, one-sided Wilcoxon test) and most of the points are above the diagonal line. We also show the random forest scatter plots for mean representations in Fig. S10b, which does not show significantly larger random forest errors compared to *β*-TCVAE (*p >* 0.05, one-sided Wilcoxon test) and might be due to the remaining correlations among mean representations of *β*-TCVAE [49]. Thus, MichiGAN with sampled representations is able to more accurately make predictions from latent space arithmetic than *β*-TCVAE. However, some of the six predictions for the missing combinations show low random forest errors from both methods, and some of the predictions from MichiGAN are only marginally better than those of *β*-TCVAE.

**Fig. 4:**
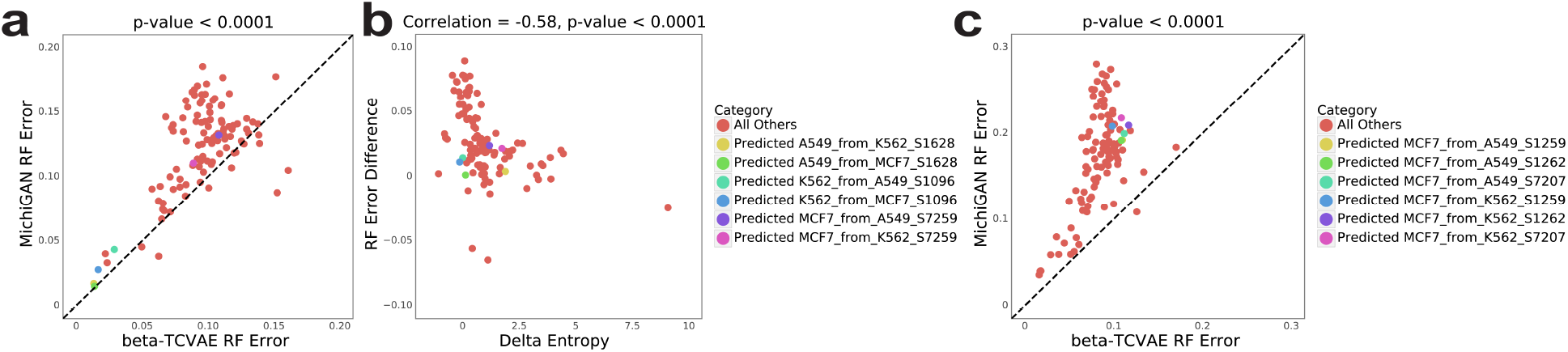
Predicting single-cell data with a perturbation by MichiGAN. The MichiGAN frameworks using sampled representations are presented. **a** Random forest (RF) errors between MichiGAN and *β*-TCVAE for all combinations trained on the large screen sci-Plex data without three combinations of A549-S1628, K562-S1096 and MCF7-S7259. **b** Scatter plots of random forest errors’ difference between MichiGAN and *β*-TCVAE versus *ΔH* trained on the large screen sci-Plex data from a. **c** Random forest errors for MichiGAN and *β*-TCVAE after selecting held-out combinations with low *ΔH*.

### 3.6 Accuracy of latent space arithmetic influences MichiGAN prediction accuracy

We next examined factors influencing the accuracy of MichiGAN predictions from latent space arithmetic. We suspected that the prediction accuracy might depend on the accuracy of the latent coordinates calculated by latent space arithmetic, which could vary depending, for example, on whether the drug exerts a consistent effect across cell types.

The quantity *ΔH* measures how accurately latent space arithmetic predicts the latent values for the held-out data. Thus, we expect that MichiGAN should be able to more accurately predict drug/cell type combinations with a small *ΔH*.

As Fig. 4b and Fig. S11a show, Δ*H* is significantly correlated with the difference in random forest error between MichiGAN and *β*-TCVAE, when sampling from either the posterior distribution of the latent representations or the posterior means. This supports our hypothesis that accuracy of the latent space arithmetic influences MichiGAN performance. To further test this, we selected the three drug/cell type combinations with the lowest overall *ΔH* values, and re-trained the network using all combinations except these three. Fig. 5 shows the predicted, real and control cells for the six predictions of the three new missing combinations based on MichiGAN using sampled representations. The predicted cells (green) overlap most parts of the real cells (blue) for all six predictions. As expected, MichiGAN predicted each of these low *ΔH* held-out combinations significantly more accurately than *β*-TCVAE (Fig. 4c).

**Fig. 5:**
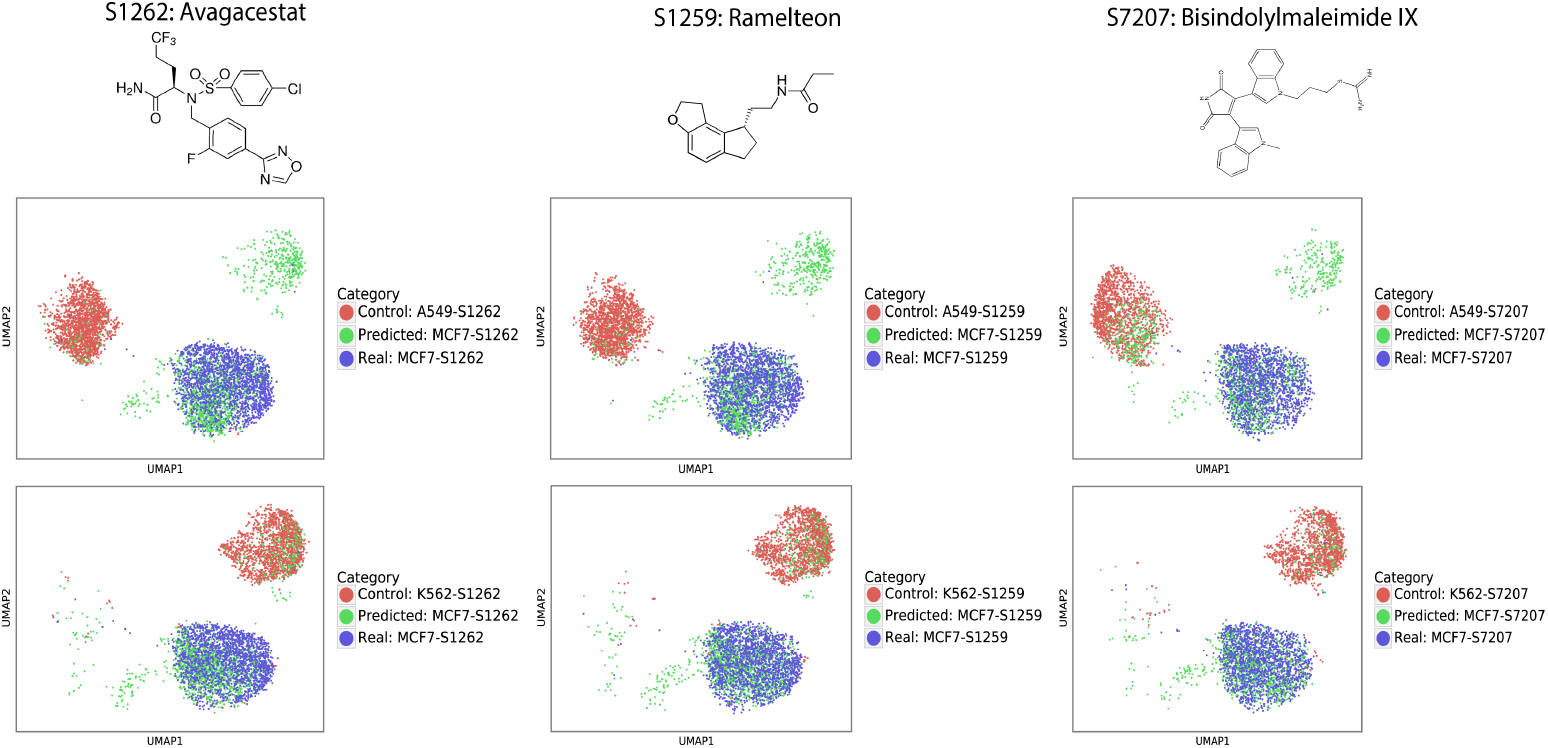
UMAP plots of the predicted (green), real (blue) and control (red) cells for 6 predictions of the three missing combinations of MCF7-S1262, MCF7-S1259 and MCF7-S7207.

## 4 Discussion

Our work provides fundamental evaluations of disentanglement performances of deep generative models on single-cell RNA-seq data. We show that combining GANs and VAEs can provide strong performance in terms of both data generation and disentanglement. The MichiGAN network provides an alternative to the current disentanglement learning literature, which focuses on learning disentangled representations through improved VAE-based or GAN-based methods, but rarely by combining them. Additionally, as the state-of-the-art in disentangled representation advances, we can immediately incorporate new approaches in the MichiGAN framework, since the training of representation and GAN are completely separate.

We envision several exciting future directions. First, it would be interesting to investigate the representations learned by *β*-VAE or *β*-TCVAE across a range of biological contexts. Second, incorporating additional state-of-the-art GAN training techniques may further improve data generation quality. Additionally, there are many other biological settings in which predicting unseen combinations of latent variables may be helpful, such as cross-species analysis or disease state prediction.

## A Appendix

The supplementary material contains two sections. Section A.1 reviews additional methods. Section A.2 gives additional experimental results.

### A.1 Additional methods

#### Real scRNA-seq datasets

The Tabula Muris dataset is a compendium of single-cell transcriptomic data from the model organism Mus musculus [13]. We processed the Tabula Muris data using SCANPY [75] and the dataset contains 41, 965 cells and 4, 062 genes from 64 cell types. The sci-Plex dataset has three cell types treated with 188 molecules targeting 22 pathways. [68]. We selected the 18 common pathways among the three cell types and chose the drug treatment from each pathway with largest number of cells. We also use SCANPY to process the data and then have 64, 050 cells and 4, 295 genes.

#### Deep generative models

The most two widely used types of deep generative models are variational autoencoders (VAEs) [41] and generative adversarial networks (GANs).

##### Variational autoencoders (VAE)

VAE has an encoder network with parameters (*ϕ*), which maps the input data (*X*) to a latent space *Z*, and a decoder network parameterized by (*θ*), which reconstructs the highdimensional data from the latent space.

Rather than learning a deterministic function for the encoder as in a conventional autoencoder, a VAE learns the mean and variance parameters of the posterior distribution over the latent variables. However, even using a factorized Gaussian prior, the posterior is intractable. Thus, VAEs perform parameter inference using variational Bayes. Following a standard mean-field approximation, one can derive an evidence lower bound (ELBO). The objective function of VAE is to maximize the ELBO or minimize its opposite with respect to *ϕ* and *θ*:

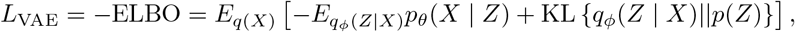

The ELBO has a nice interpretation: the first term is reconstruction error and the second term is the Kullback-Leibler divergence between the posterior and prior distributions of the latent variables (*Z*). If the prior distribution *p*(*Z*) is factorized Gaussian or uniform distribution, the KL divergence encourages the latent factors to be statistically independent, which may contribute to the good disentanglement performance of VAEs. This effect can be further enhanced by introducing a weight *β* to place more emphasis on the KL divergence at the cost of reconstruction error, an approach called *β*-VAE [32].

*β-TCVAE* The total correlation variational autoencoder (*β*-TCVAE) is a VAE extension that further promotes disentanglement. The KL divergence of VAE can be further decomposed into several parts:

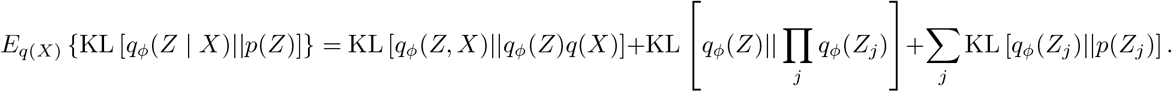

The first part is referred to as the index-code mutual information (MI), the second part is the total correlation (TC) and the third part is the dimension-wise KL divergence [11]. The total correlation is the most important term for learning disentangled representations, while penalizing the two other parts does not directly improve the disentanglement performance and increases the reconstruction error.

The *β*-TCVAE specifically penalizes the TC in the loss function [11,40]:

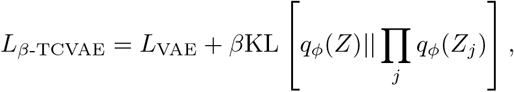

where *β* = 0 gives the VAE loss function. There is no closed form for the total correlation of the latent representation, so *β*-TCVAE approximates it as follows:

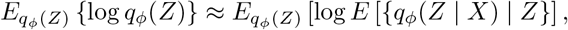

and

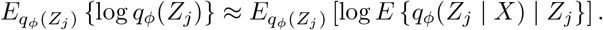

Estimating TC is difficult from a small minibatch, so we utilize the minibatch stratified sampling mentioned in [11] to estimate *E*{*q_ϕ_*(*Z* | *X*) | *Z*} during training.

##### Generative adversarial networks (GANs)

A generative adversarial network consists of a generator network *G* and a discriminator network *D*. There are many types of GANs, but we specifically focus on Wasserstein GAN with gradient penalty (WGAN-GP) [28], which significantly stabilizes GAN training. The discriminator loss function for WGAN-GP is

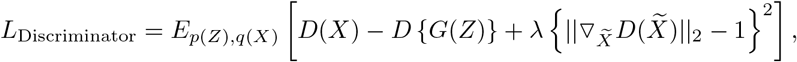

where ∇*_X_D* is the gradient of the discriminator on input *X* and 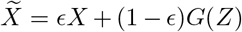 with *ϵ* sampled from a uniform distribution on [0,1]. The generator loss function for WGAN-GP is

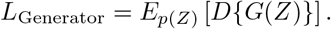

Upon convergence, WGAN-GP gives the generated data distribution *G*(*Z*) that matches the real data distribution *p*(*X*).

##### InfoGAN and ss-InfoGAN

The Information Maximizing Generative Adversarial Networks (InfoGAN) framework extends the regular GAN to encourage disentanglement [12]. The InfoGAN decomposes the latent variables into latent code *C* and noise *Z*. To encourage disentanglement, InfoGAN maximizes the mutual information between the latent code and the generated data. To estimate mutual information, InfoGAN relies on an additional network Q that takes generated data as input and predicts the code *Q*(*C* | *X*) that generated the data. *Q*(*C* | *X*) is very similar to an encoder in a VAE and estimates a posterior distribution in the same family as the prior distribution of the code *p*(*C*). InfoGAN then maximizes mutual information between the code and generated data with the following loss functions for the discriminator and generator:

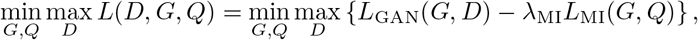

where 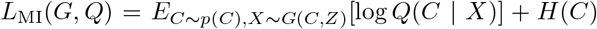 is a lower bound for the mutual information between *C* and *X* and *H*(*C*) is the entropy of the codes. We implemented InfoGAN with the Wasserstein distance, which we refer to as InfoWGAN-GP. We choose a factorized normal distribution with unit variance for *Q*(*C* | *X*) (the unit variance stabilizes InfoGAN training [12, 46]).

InfoGAN architecture can also be extended to semi-supervised InfoGAN (ssInfoGAN), if labels are available for some or all of the data points [67]. The ssInfoGAN maximizes mutual information not only between the generated data and the codes, but also between the real data and corresponding labels. This guides the learned codes to reflect the label information.

##### Conditional GAN and PCGAN

The conditional GAN extends GANs to respect the relationship between generated data and known labels [54]. There are many different network architectures for conditional GAN [54, 62, 56], but we found that the conditional GAN with projection discriminator (PCGAN) [55] works best. A recent paper similarly found that PCGAN worked well for single-cell RNA-seq data [51]. The original PCGAN paper mentions that the projection discriminator works most effectively when the conditional distribution *p*(*C*|*X*) is unimodal. One theoretical reason why PCGAN may be well-suited for MichiGAN is that the posterior multivariate Gaussian distributions of latent variables from VAEs are, in fact, unimodal.

In implementing the PCGAN, we do not use the conditional batch normalization or spectral normalization mentioned in [55], but instead use standard batch normalization and Wasserstein GAN with gradient penalty. Thus we refer to this approach as PCWGAN-GP.

#### Mutual information

Following [11], we measure the disentanglement performance of the representations using mutual information gap (MIG). Denote *p*(*V_k_*) and *p*(*X* | *V_k_*) as the probability of a ground-truth variable *V_k_* and the conditional probability of the data *X* under *V_k_*. Given *q_ϕ_*(*Z_j_*, *V_k_*) = ∫*_X_ p*(*V_k_*)*p*(*X* | *V_k_*)*q_ϕ_*(*Z_j_* | *X*)*dX*, the mutual information between a latent variable *Z_j_* and a ground-truth variable *V_k_* is

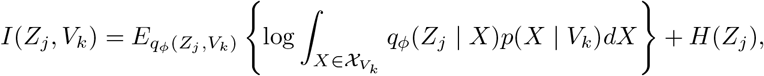

defined as
where 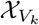 is the support of *X* ~ *p*(*X* | *V_k_*) and *H*(*Z_j_*) is the entropy of *Z_j_*. Due to the different variabilities of the ground-truth variables, the normalized mutual information is better to be used with a normalization term of *H*(*V_k_*), the entropy of *V_k_*. The posterior distribution *q_ϕ_*(*Z_j_* | *X*) is obtained from the encoder (for VAEs) or the derived posterior distribution for probabilistic PCA [8]. With *K* ground truth variables {*V*_1_, …, *V_k_*}, the mutual information gap (MIG) is further defined as

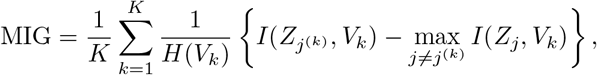

where *j*^(*k*)^ = arg max*_j_ I*(*Z_j_*, *V_k_*).

The MIG metric is taking the difference between largest and the second largest normalized mutual information values for each ground-truth variable and compute the average across all ground-truth variables, indicating how much a unique representation can outperform other representations on representing each variable and quantifies the disentanglement performance. If the averaged gap is high, more mutual information is restored in one specific representation for each ground-truth variable. As described in [11], the MIG metric has the axis-alignment property and is unbiased for all hyperparameter settings and generally applicable.

#### Implementation

The VAE-based methods use multilayer perceptron (MLP) and have two fully-connected (FC) hidden layers with 512 and 256 neurons, followed by separate parameters for mean and variance of the latent representation. The first two hidden layers in the decoder have 256 and 512 neurons, while the last layer gives mean gene expression and has the same number of neurons as the number of genes. Each hidden layer utilizes batch normalization, activated by Rectified Linear Unit (ReLU) or Leaky ReLU. Each hidden layer employs dropout regularization, with a dropout probability of 0.2. We also experimented with three hidden layers for the VAE encoders, but found that the training became unstable. This is consistent with a previous report [34] that found most VAEs for biological data have only two hidden layers. The GAN-based methods also use MLP for both generator and discriminator. There are three FC hidden layers with 256, 512 and 1024 neurons as well as three hidden layers with 1024, 512 and 10 neurons from data to output. The hidden layers of GANs also have Batch Normalization and ReLU or Leaky ReLU activation. The generator uses dropout regularization with dropout probability of 0.2 for each hidden layer. The VAE-based methods are trained with Adam optimization, while the GAN-based methods are trained with Adam and the gradient prediction method [77]. The batch sizes of VAE-based methods and GAN-based methods are 128 and 32, respectively. All the hyperparameters of each method on different datasets are tuned for the optimal results.

We trained all models for 1,000 epochs and used 10 latent variables. We used *β* = 50 and *β* = 10 for *β*-TCVAE on the simulated single-cell RNA-seq datasets with non-linear and linear step variable, respectively. For the two real scRNA-seq datasets, we used *β* = 100. We used 118-dimensional Gaussian noise for MichiGAN. We also trained an additional WGAN-GP model with 128-dimensional Gaussian noise for comparison of latent traversals. All models were implemented in TensorFlow.

### A.2 Additional results

#### Variational autoencoders learn disentangled representations of single-cell data

We present Fig. S1, Fig. S2 and Fig. S3 to compare the disentangled representations of different deep generative models on single-cell data.

**Fig. S1:**
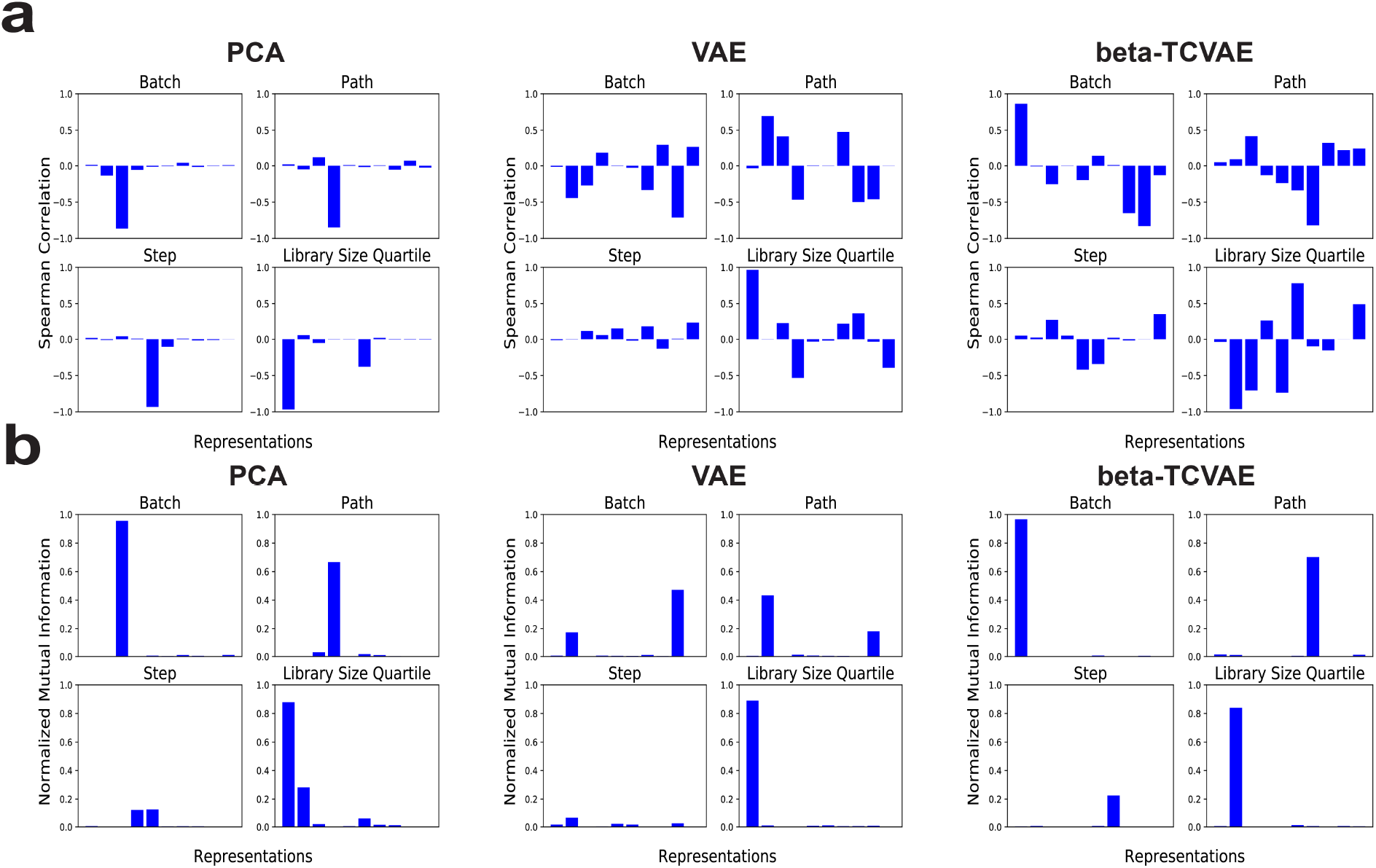
Learned representations of simulated single-cell data with non-linear step. **a** Bar plots of Spearman correlations between ten representations and each of the four ground-truth variables for PCA, VAE and *β*-TCVAE. **b** Bar plots of normalized mutual information between ten representations and each of the four ground-truth variables for PCA, VAE and *β*-TCVAE.

**Fig. S2:**
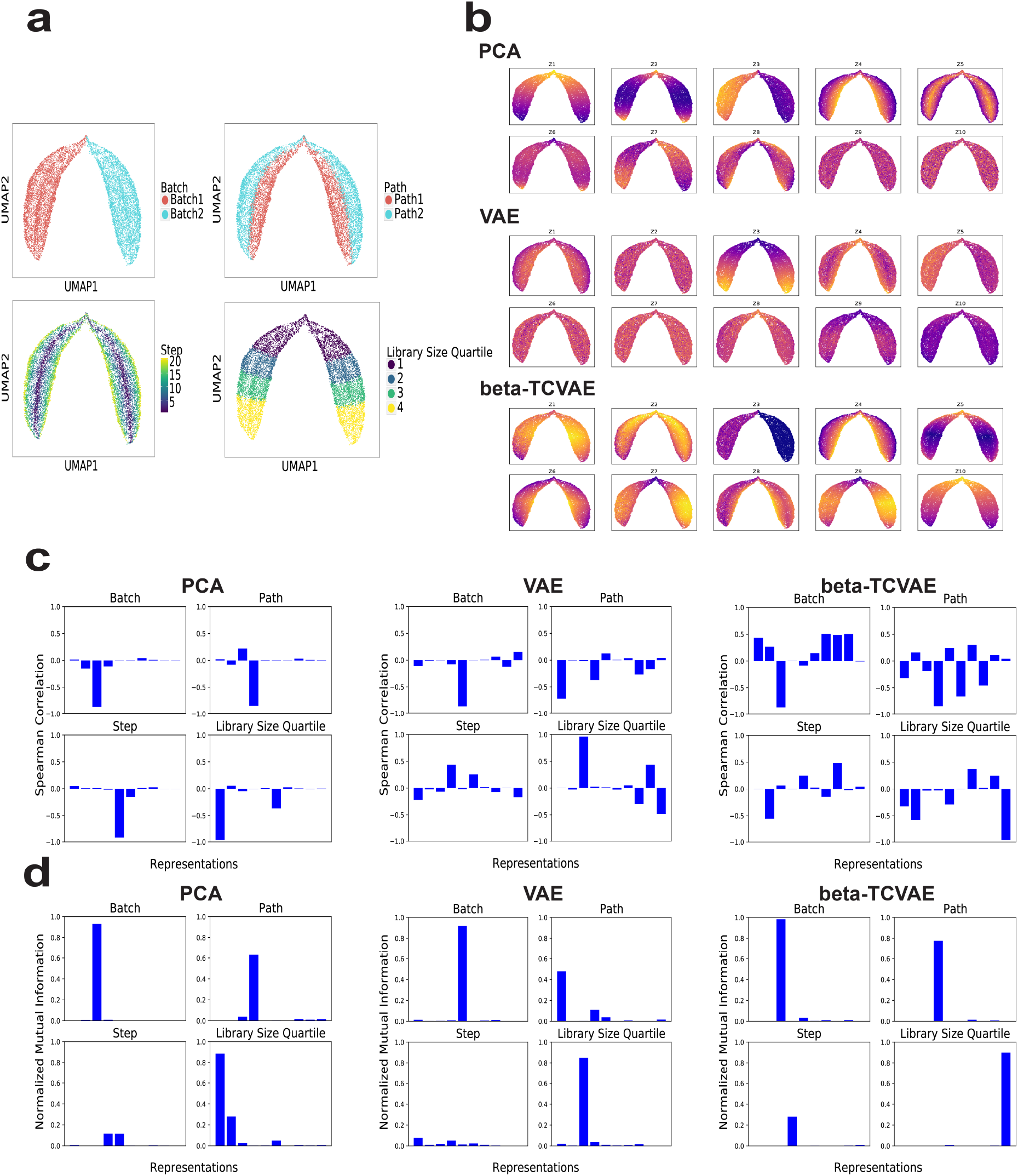
Learned representations of simulated single-cell data with linear step. **a** UMAP plots of data by batch, path, step and library size quartile. **b** UMAP plots of data colored by the ten representations learned by PCA, VAE and *β*-TCVAE. **c** Bar plots of Spearman correlations between ten representations and each of the four ground-truth variables for PCA, VAE and *β*-TCVAE. **d** Bar plots of normalized mutual information between ten representations and each of the four ground-truth variables for PCA, VAE and *β*-TCVAE

**Fig. S3:**
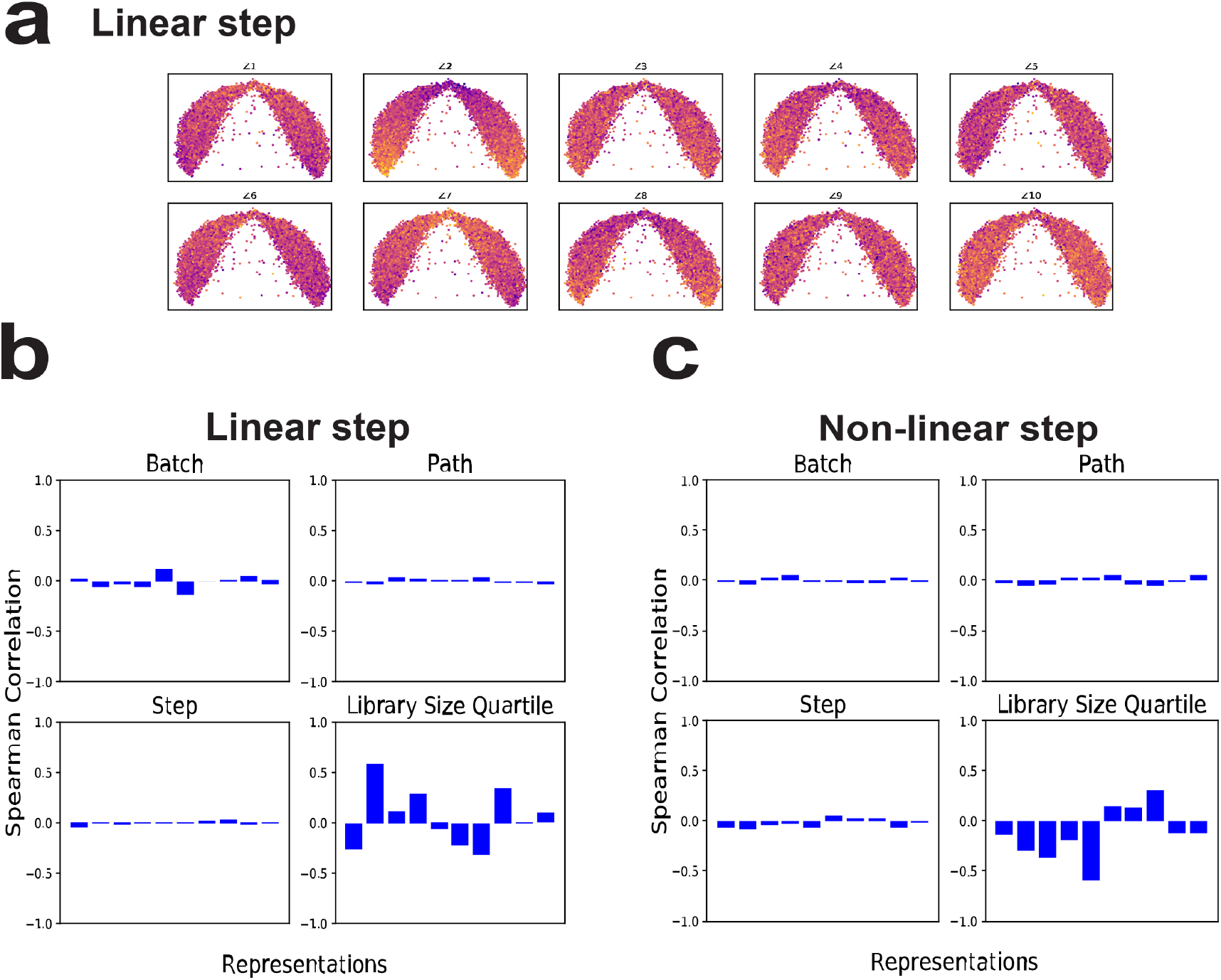
Learned representations of InfoWGAN-GP for the simulated single-cell data. **a** UMAP plots of the simulated data with linear step colored by the ten representations learned by InfoWGAN-GP. **b** Bar plots of Spearman correlations between ten representations and each of the four ground-truth variables for InfoWGAN-GP on the simulated data with linear step. **c** Bar plots of Spearman correlations between ten representations and each of the four ground-truth variables for InfoWGAN-GP on the simulated data with non-linear step.

#### GANs generate more realistic single-cell expression profiles than VAEs

We present Fig. S4 to compare the generation performance of different deep generative models on single-cell data.

**Fig. S4:**
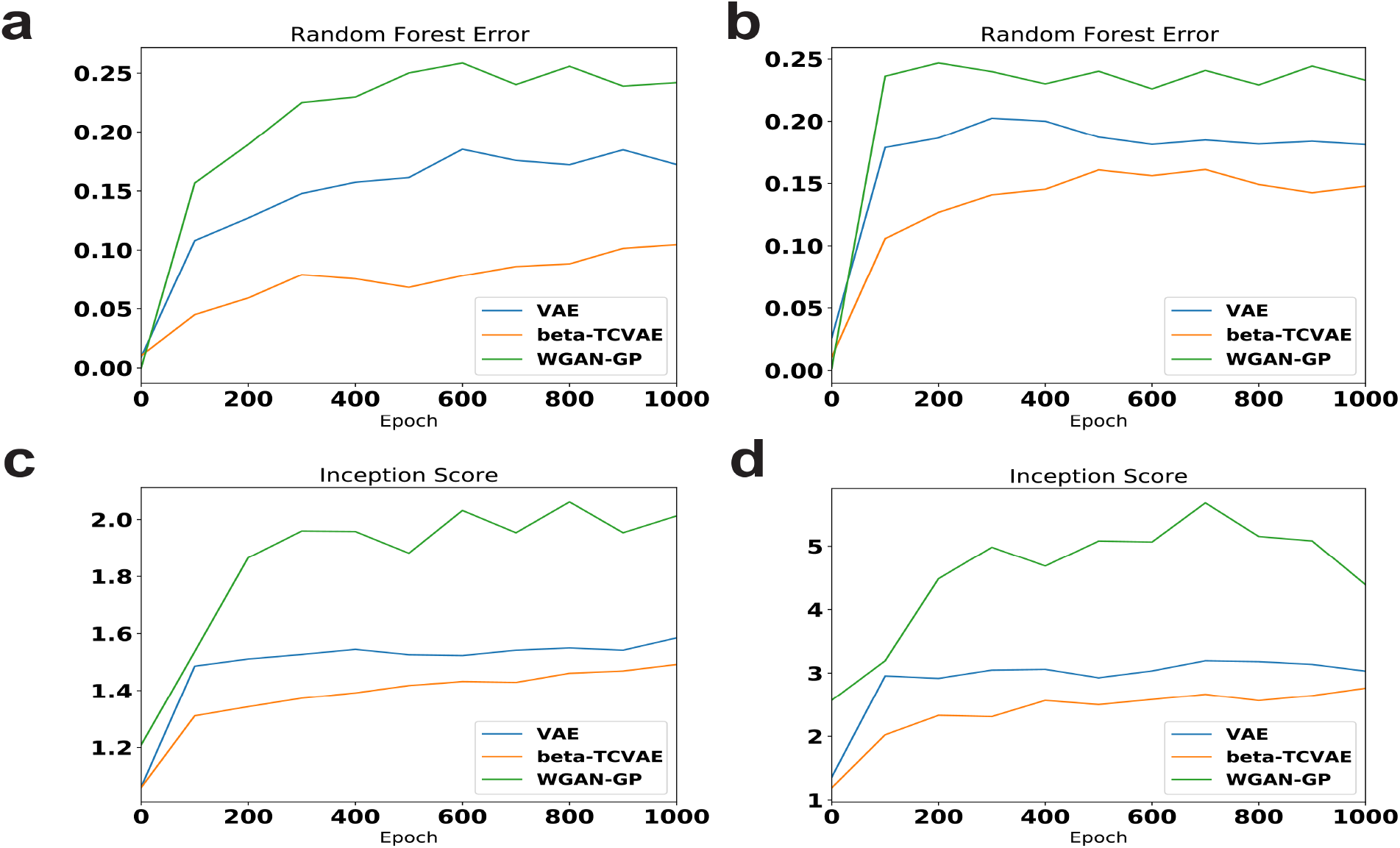
Generation performances of VAE, *β*-TCVAE ad WGAN-GP on the Tabula Muris heart data and the whole Tabula Muris data. **a** Random forest error of the three methods on the Tabula Muris heart data during training. **b** Random forest error of the three methods on the whole Tabula Muris data during training. **c** Inception score of the three methods on the Tabula Muris heart data during training. **d** Inception score of the three methods on the whole Tabula Muris data during training.

#### MichiGAN samples from disentangled representations without sacrificing generation performance

We present Fig. S5 and Fig. S6 to show that MichiGAN samples from disentangled representations without sacrificing generation performance.

**Fig. S5:**
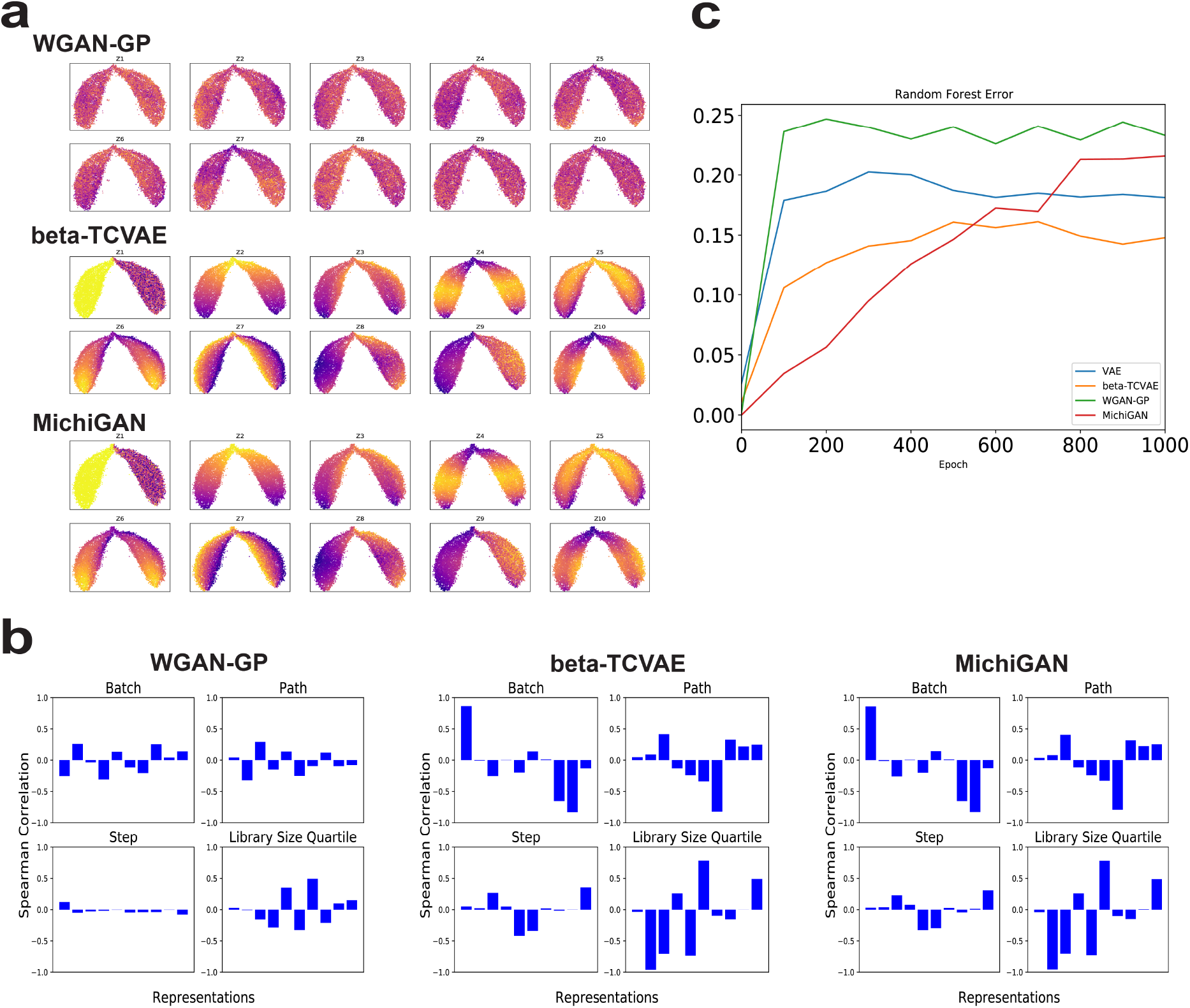
Disentanglement and generation performance of WGAN-GP, *β*-TCVAE and MichiGAN. **a** UMAP plots of generated data colored by the ten representations of WGAN-GP, *β*-TCVAE and MichiGAN on the simulated data with non-linear step. **b** Bar plots of Spearman correlations between ten representations and each of the four ground-truth or inferred variables for WGAN, *β*-TCVAE and MichiGAN on the simulated data with non-linear step. **c** Random forest error of VAE, *β*-TCVAE, WGAN-GP and MichiGAN on the whole Tabula Muris data during training.

**Fig. S6:**
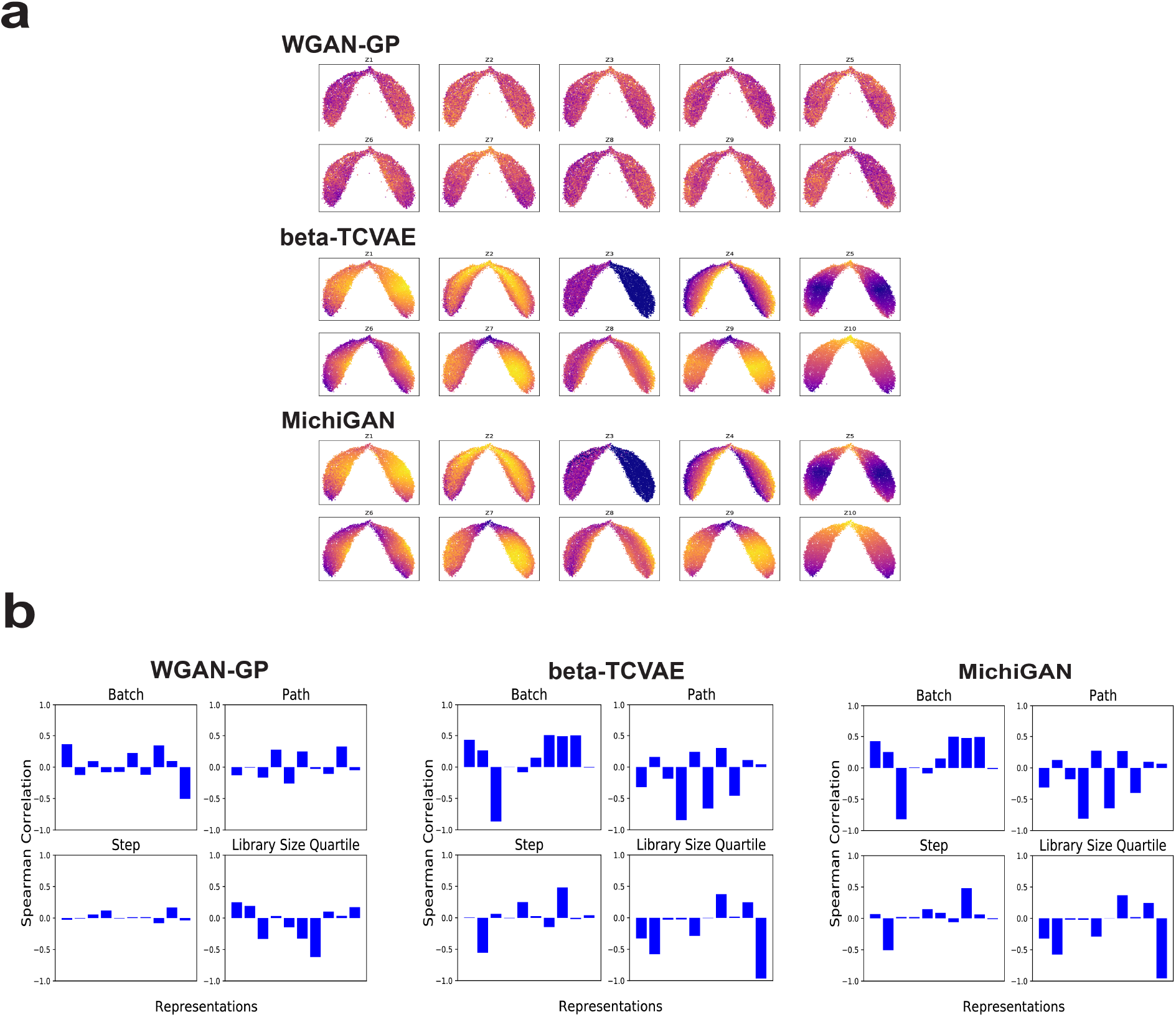
Disentanglement and generation performance of WGAN-GP, *β*-TCVAE and MichiGAN. **a** UMAP plots of generated data colored by the ten representations of WGAN-GP, *β*-TCVAE and MichiGAN on the simulated data with linear step. **b** Bar plots of Spearman correlations between ten representations and each of the four ground-truth or inferred variables for WGAN, *β*-TCVAE and MichiGAN on the simulated data with linear step.

#### MichiGAN enables semantically meaningful latent traversals

We present Fig. S7, Fig. S8 and Fig. S9 to show that MichiGAN enables semantically meaningful latent traversals, while WGAN-GP is entangled.

**Fig. S7:**
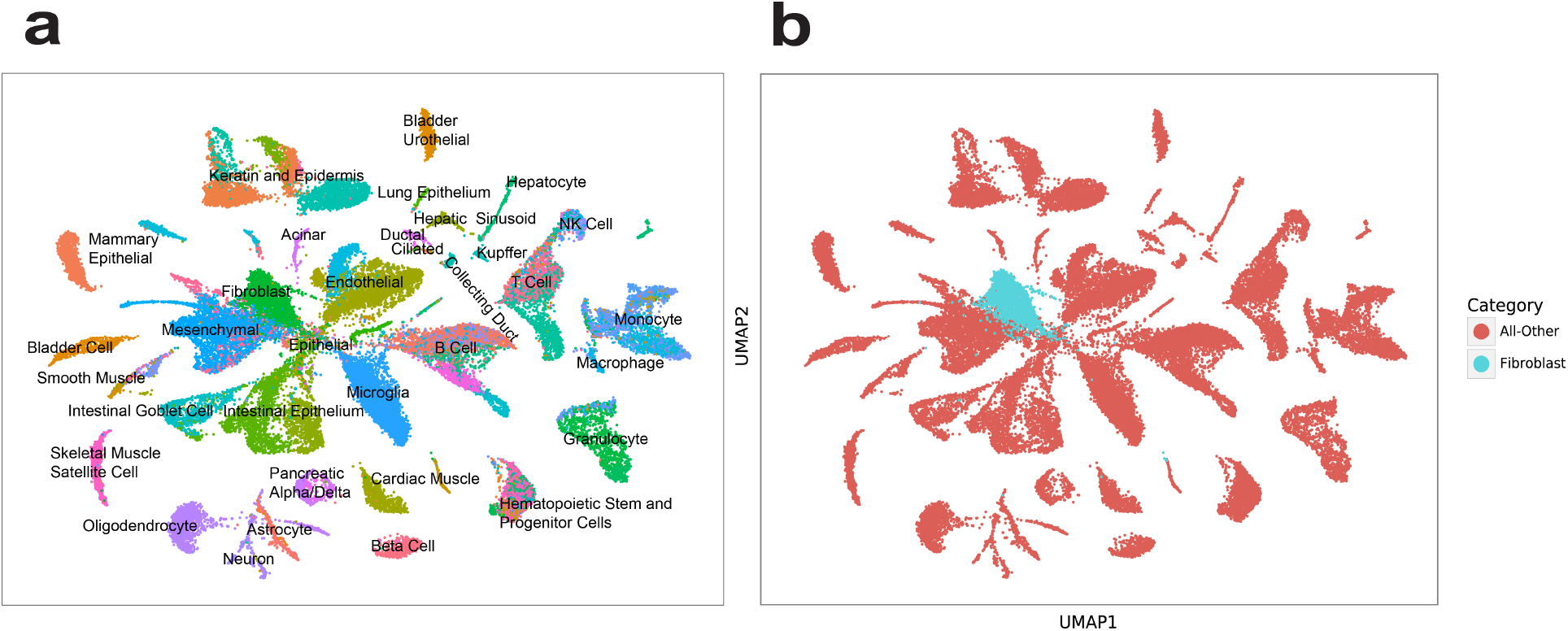
The whole Tabula Muris data. **a** UMAP plot of the whole Tabula Muris data colored by cell type. **b** UMAP plot of the 2026 fibroblast cells in heart within the whole Tabula Muris data.

**Fig. S8:**
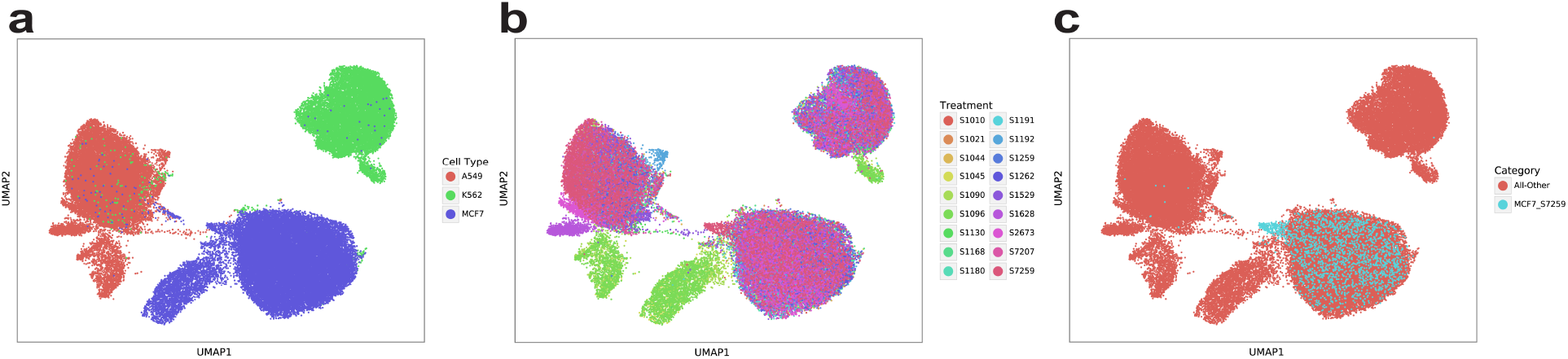
The large screen sci-Plex data. **a** UMAP plot of the sci-Plex data colored by cell type. **b** UMAP plot of the sci-Plex data colored by drug treatment. **c** UMAP plot of the 2014 cells with MCF7 cell type and S7259 treatment within the sci-Plex data.

**Fig. S9:**
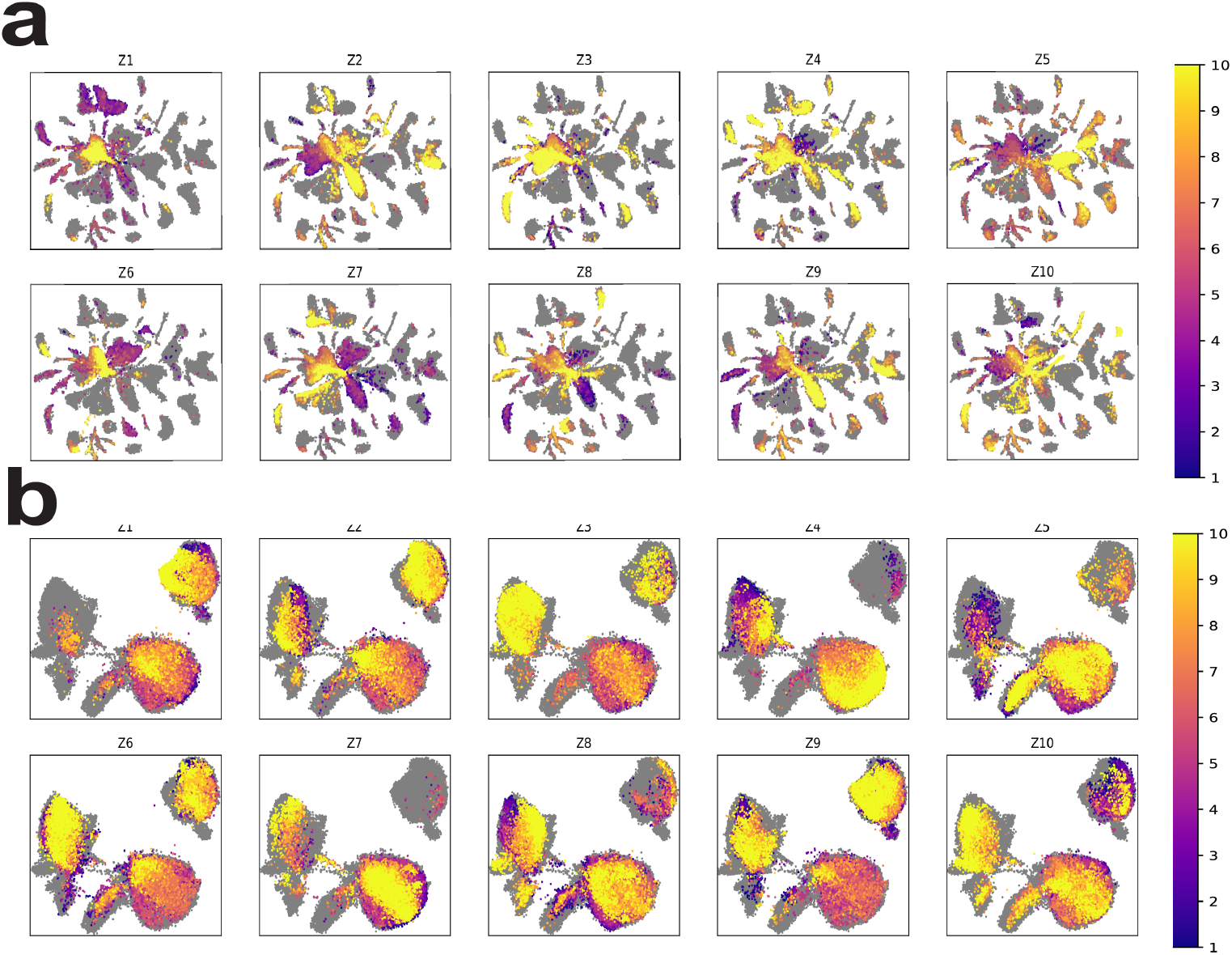
UMAP plots of generated data via latent traversals. **a** UMAP plot of latent traversals of the 10 representations of latent values that generate data closest to fibroblast cells in heart within the Tabula Muris data using WGAN-GP with 10 dimensions. **b** UMAP plot of latent traversals of the 10 representations of latent values that generate data closest to MCF7-S7259 cells within the sci-Plex data using WGAN-GP with 10 dimensions.

#### MichiGAN predicts single-cell gene expression changes under unseen drug treatments

We show the information of large screen sci-Plex data in Table S1. We present Fig. S10 to show that MichiGAN predicts single-cell gene expression changes under unseen drug treatments.

**Table S1:**
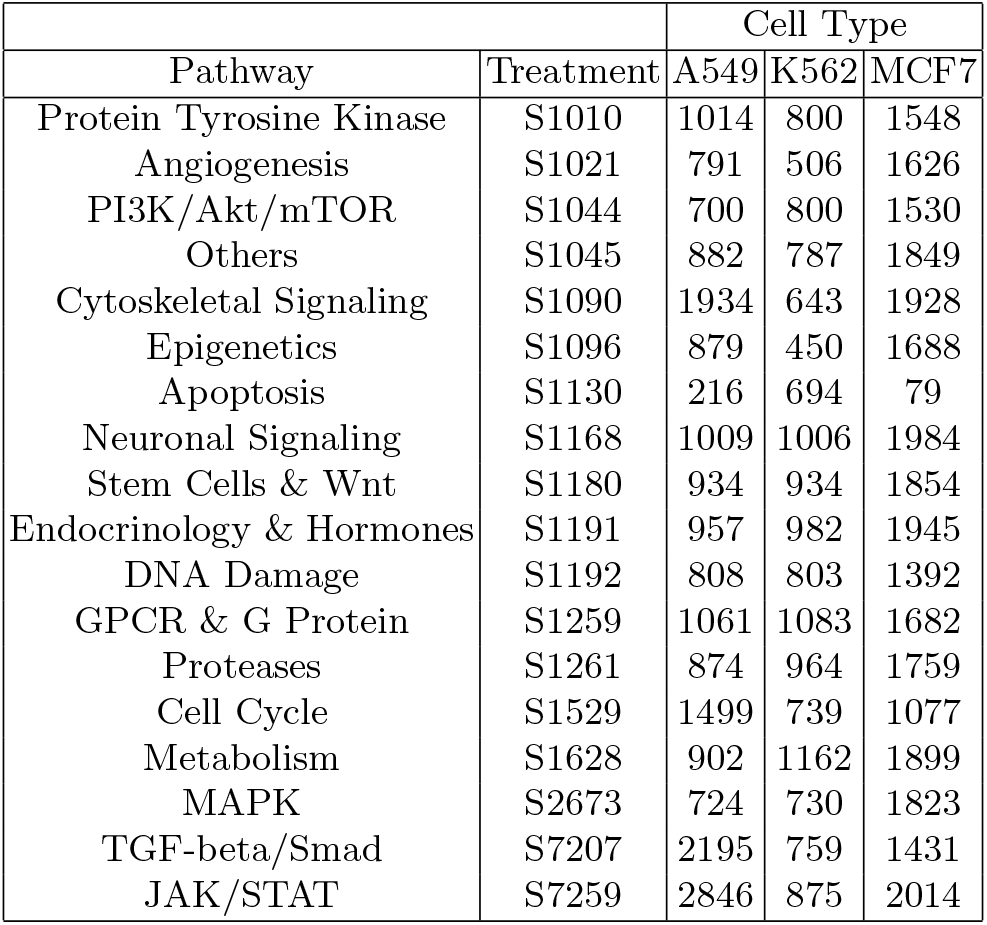
Numbers of cells of three cell types and 18 pathways or treatments of subsampled sci-Plex data

| Pathway | Treatment | Cell Type |  |  |
| --- | --- | --- | --- | --- |
|  |  | A549 | K562 | MCF7 |
| Protein Tyrosine Kinase | S1010 | 1014 | 800 | 1548 |
| Angiogenesis | S1021 | 791 | 506 | 1626 |
| PI3K/Akt/mTOR | S1044 | 700 | 800 | 1530 |
| Others | S1045 | 882 | 787 | 1849 |
| Cytoskeletal Signaling | S1090 | 1934 | 643 | 1928 |
| Epigenetics | S1096 | 879 | 450 | 1688 |
| Apoptosis | S1130 | 216 | 694 | 79 |
| Neuronal Signaling | S1168 | 1009 | 1006 | 1984 |
| Stem Cells & Wnt | S1180 | 934 | 934 | 1854 |
| Endocrinology & Hormones | S1191 | 957 | 982 | 1945 |
| DNA Damage | S1192 | 808 | 803 | 1392 |
| GPCR & G Protein | S1259 | 1061 | 1083 | 1682 |
| Proteases | S1261 | 874 | 964 | 1759 |
| Cell Cycle | S1529 | 1499 | 739 | 1077 |
| Metabolism | S1628 | 902 | 1162 | 1899 |
| MAPK | S2673 | 724 | 730 | 1823 |
| TGF-beta/Smad | S7207 | 2195 | 759 | 1431 |
| JAK/STAT | S7259 | 2846 | 875 | 2014 |

**Fig. S10:**
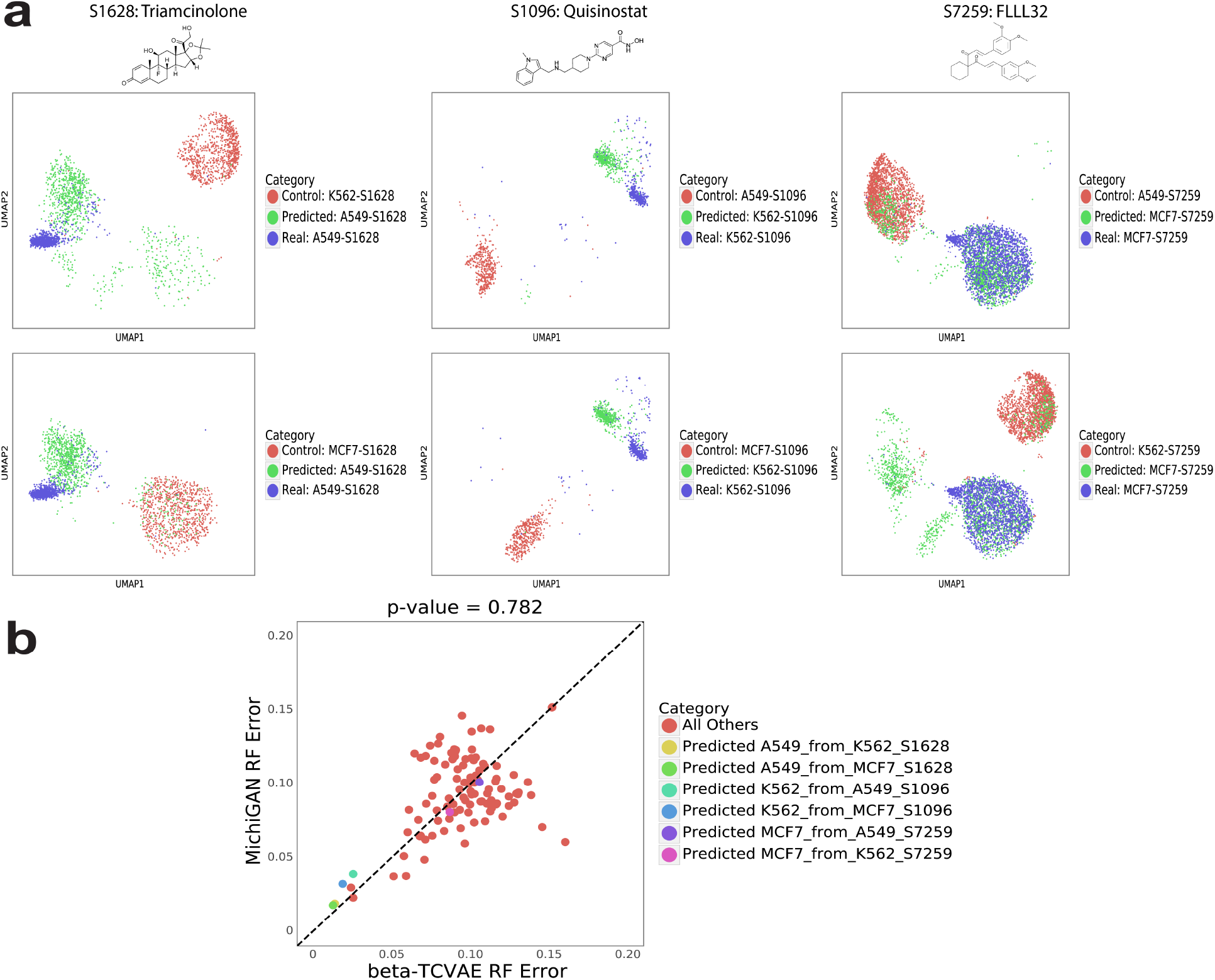
Performance of MichiGAN trained from *β*-TCVAE mean representations on sci-Plex data. **a** UMAP plots of the predicted (green), real (blue) and control (red) cells for 6 predictions of the three missing combinations of A549-S1628, K562-S1096 and MCF7-S7259. **b** Random forest errors between MichiGAN and *β*-TCVAE for all combinations.

#### Accuracy of latent space arithmetic influences MichiGAN prediction accuracy

We present Fig. S11 to show that accuracy of latent space arithmetic influences MichiGAN prediction accuracy.

**Fig. S11:**
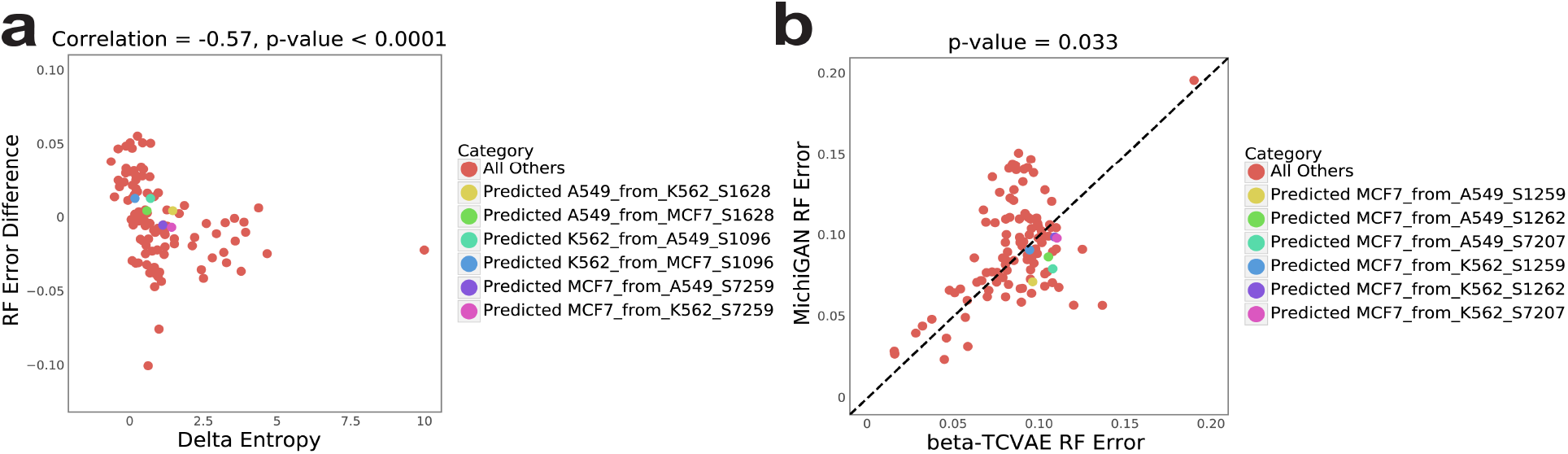
Performance of MichiGAN trained from *β*-TCVAE mean representations on held-out combinations selected for low delta entropy. **a** Scatter plots of random forest errors’ difference between MichiGAN and *β*-TCVAE versus delta entropy for MichiGAN trained with mean representations on the large screen sci-Plex data without three combinations of A549-S1628, K562-S1096 and MCF7-S7259. **b** Random forest error differences between MichiGAN and *β*-TCVAE after selecting held-out combinations with low *ΔH*.

## Footnotes

1 Code is available at: https://github.com/welch-lab/MichiGAN.

